# Intrinsic neural timescales attenuate information transfer along the uni-transmodal hierarchy

**DOI:** 10.1101/2023.07.28.551047

**Authors:** Yasir Çatal, Angelika Wolman, Samira Abbasi, Georg Northoff

## Abstract

The brain’s intrinsic timescales are organized in a hierarchy with shorter timescales in sensory regions and longer ones in associative regions. This timescale hierarchy overlaps with the timing demands of sensory information. Our question was how does this timescale hierarchy affect information transfer. We used a model of the timescale hierarchy based on connected excitatory and inhibitory populations across the cortex. We found that a hierarchy of information transfer follows the hierarchy of timescales with higher information transfer in sensory areas while it is lower in associative regions. Probing the effect of changes in timescale hierarchy on information transfer, we changed various model parameters which all, through, the loss of hierarchy, induced increased information transfer. Finally, the steepness of the timescale hierarchy relates negatively to total information transfer. Human MEG data confirmed our results. In sum, we demonstrate a key role of the brain’s timescale hierarchy in mediating information transfer.

## Introduction

The brain continuously processes information with a variety of timescales ranging from milliseconds (flashes of light, syllables) to minutes (watching a movie, a conversation). To make sense of this sensory bombardment, the brain has to possess certain biocomputational mechanisms to modulate and, to a certain degree, attenuate the seemingly overwhelming information transfer. Recent studies point out that the brain possesses different timescales in its spontaneous activity, called intrinsic neural timescales (INTs)^1–3^. Different timescales of neural activity are situated on a rostrocaudal axis of increasing synaptic excitation^4^ and increasing inhibition which gates long-range excitation as opposed to gating output of that region^5,6^. Correspondingly, sensory regions show fast timescales in their activity which is suitable for processing the continuously changing stimuli while transmodal areas implicated in higher-order processes exhibit longer timescales, suited well for integrating information from different modalities ^3,7,8^. This has been observed in a variety of imaging modalities including single neuron recordings^8^, ECoG^9^, fMRI^10–13^, EEG^14,15^ and MEG^16^; in resting state^11,13,16^ as well as in various task states^8,14,17^. Such occurrence across different levels and scales suggests a key role and function of the timescale hierarchy in information processing. Does the hierarchy of timescales in the brain’s neural activity modulate its information transfer from uni- to transmodal regions? Addressing this gap in our knowledge is the goal of our study.

In a series of impressive studies, the group around Hasson directly related the brain’s timescales to the timescales of the external stimuli such as narratives^18^ and music^19^. These studies show that the brain’s different regions’ so-called temporal receptive windows (TRW) are sensitive to the stimuli that matches their own intrinsic timescales^3,20^. Such correspondence of the timescales of both brain and stimuli raises the question whether the brain’s information transfer is also organized along its uni-transmodal timescale hierarchy. This remains an open question, though. How and in what way do the timescale hierarchy modulate and transfer the incoming information across its various regions? Does the information transfer change with the hierarchy of timescales along their uni-transmodal gradient?

Addressing these questions, we used a computational model and manipulated its different parameters (excitation-inhibition balance, intrinsic time constants etc.) while, at the same time, measuring its information transfer from uni- to transmodal regions. This was complemented by empirical data, e.g., MEG, to support the assumed relationship of the timescale hierarchy with a corresponding uni-transmodal gradient in information transfer.

The mathematical model we used (described in^21^) is based on one excitatory and one inhibitory population in 29 regions in the cortex (figure 1C). Excitatory and inhibitory populations are connected to each other via local connections whereas different regions in the brain are connected via long range excitatory coupling *μ* that scales the structural connections that were obtained from macaque data^22^. In accordance with empirical observations^4^, synaptic excitation gradually increases over the shift from uni- to transmodal regions in the model. The steepness of this uni-transmodal increase is controlled by *η* The dynamics of firing rates is controlled by a linear-threshold rectifier function whose slope is parametrized by *β* controls Finally, the time constant τ how fast the dynamics evolve.

**Figure 1.**
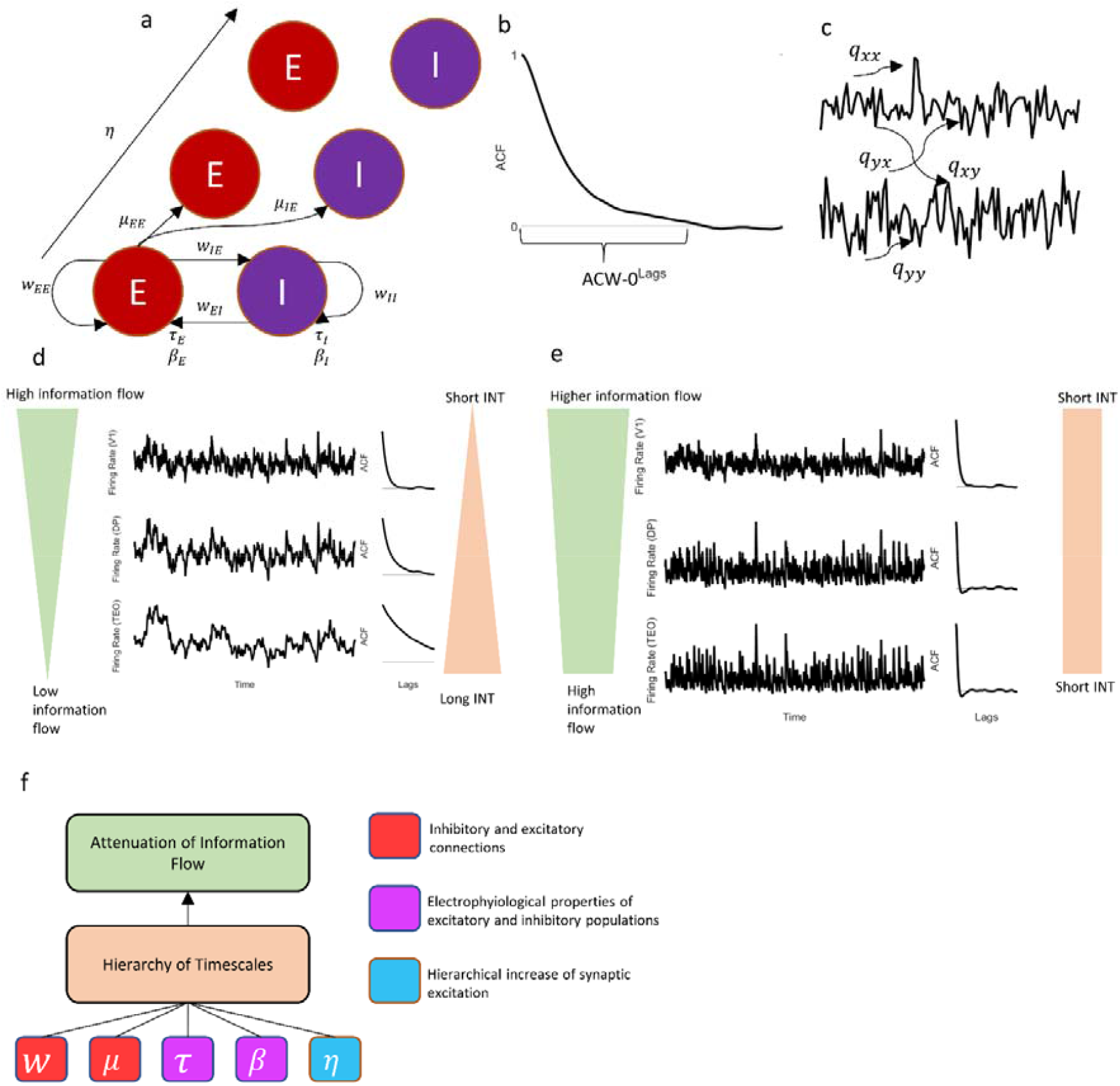
Overview of the paper. A. ACW-0 is calculated as the lag where autocorrelation function touches 0. It is used as an index of intrinsic neural timescales (INT). B. Transfer entropy is a directed measure of effective connectivity. The transfer entropy from y to x is calculated as how much our uncertainty reduces about the future of x if we know the past q_yx values of y in addition to past q_xx values of x .We used a data-driven method to determine the Markov lags (q values) in the calculation. C. Model architecture. E and I denote excitatory and inhibitory populations. For other parameters, see methods. D. A hierarchy of information transfer follows the hierarchy of INTs. Sensory regions have shorter INT and higher information transfer, and vice versa for associative regions. E. When the timescale hierarchy is lost, total information transfer increases. F. Proposed model that describes the relationship between the timescale hierarchy, information transfer and the mathematical model. We hypothesize that changes in model parameters disrupt the timescale hierarchy, which attenuates the information transfer from lower to higher areas. When the downsampling is lost, total transfer entropy increases. The colors orange, purple and teal relate to the what the model parameters do.

To estimate INTs, we used a parameter called autocorrelation window (ACW) which is a scalar value extracted from the autocorrelation function (ACF) of a time series^8,10–12,14,15,17,23^ (figure 1A). ACF is computed as the correlation of a time series with itself on different time lags. This way, it represents the similarity of a signal with itself over time. A slowly decaying ACF would be the outcome of a signal that is similar to itself for a longer duration, changing slowly on longer timescales and vice versa. ACW is the lag where ACF crosses a threshold, for example, 0.5 (ACW-50) ^14–16^or 0 (ACW-0)^16^; as is more informative of both neural^16^ and psychological^14,15^ levels, we used the latter in our study.

The quantification of information transfer was performed on an information-theoretical framework^24^ (figure 1B). A relatively new bidirectional, nonparametric and model-free measure called transfer entropy^25–27^ (TE) was found to be useful as a measure of directed information transfer from one region to another, i.e., effective connectivity in neuroimaging data^28–31^. TE quantifies the reduction of uncertainty in the future values of time series x when the past values of another time series y in the same system is known in addition to past values in x. A high TE value means there is high information transfer from y to x while a low TE value indicates low information transfer. Due to its nonparametric nature, TE can capture nonlinear relationships that other measures based on vector autoregressive models (such as Granger causality) would miss^27^. Based on the definitions of these measures, we focused on TE and ACW to probe the relation of information transfer with INTs.

Our question is driven by the sampling theorem in the signal processing literature^40^. The sampling theorem states that the sampling rate puts an upper bound on the highest frequency of information that can be captured by a recording device. Echoing previous work by Golesorkhi et al.^2^, we hypothesized that the INT of a region serves as its sampling rate. A region that is intrinsically “slow” will not be able to encode the high-frequency information coming towards it. This will result in attenuation and downsampling of information along the timescale hierarchy (figure 1d, e, and f). We can rephrase the main goal of the study as how the hierarchical structure of sampling rates in a biological network influence the information transfer.

Our first specific aim was to demonstrate the relationship between the uni-transmodal hierarchy of INTs with information transfer. For that purpose, we focused on the visual cortical stream regions in the mathematical model described above. For this aim, we simulated the firing rates in the model by giving two kinds of input. First, we gave the early sensory area V1 a gaussian noise and small noise to all the other areas which represent their intrinsic dynamics (i.e. resting state). We calculated ACW-0 values on this state. Second, we gave added boxcar inputs (1s where there is input, 0s where not) to V1 on top of the noise to simulate a task-like state that mimics empirical MEG data (see below). We calculated TE values in this task-like state.

The second aim focused on the potentially causal impact of the uni-transmodal hierarchy on the relationship of INTs and information transfer. In a mathematical analysis of the model we used^32^ three necessary and sufficient conditions to establish the hierarchy of timescales: (i) distinct electrophysiological properties of excitatory and inhibitory populations (through *β* and *τ*); (ii) local inhibition sufficient (through *μ* and *w*) to counter and balance long-range excitation; and (iii) a hierarchical increase of synaptic excitation (through *η*). Inspired by this work, we changed different parameters in the model that enable these three conditions and investigated how these changes affect the ACW-TE relationship. Based on these necessary conditions^32^, we expected major changes in the relationship of ACW and TE when changing the model parameters related to excitation, inhibition and the steepness of the hierarchy itself.

The third specific aim consisted in replicating the relationship of INT and information transfer that we observed in the computational model using empirical data. For that purpose, we used the Human Connectome Project’s open-access MEG dataset^33–35^ and investigated the relationship of ACW and TE in the regions that were significantly recruited and activated during two tasks, namely working memory and motor. After source reconstruction, we identified the relevant regions of interest (ROI) based on prior studies^36,37^ that used the same task design and calculated ACW,TE and their relationship across the temporal (e.g., ACW-based) hierarchy across these regions.

We expected information transfer to be sensitive to timescale hierarchy. Regions higher in the hierarchy exhibit longer ACW and therefore allow for higher degrees of temporal integration of information than unimodal regions with their shorter ACW^1,14^. Concerning these regions’ information processing, this means that the transmodal regions’ longer timescales more strongly lump together and thereby integrate different information^38,39^ compared to the shorter timescales of the unimodal regions. Following that, we assumed decreased and thus attenuated information transfer in transmodal regions compared to the unimodal regions.

We found three pervasive phenomena: (i) a hierarchy of inter-regional information transfer overlaps with the hierarchy of INTs with higher information transfer in sensory areas and lower in associative ones; (ii) the total information transfer in the whole system increases when the hierarchy of timescales is flattened by manipulation of the various model parameters; (iii) the total information transfer is negatively correlated with the steepness of the INT hierarchy. Analysis of human MEG data confirmed our findings described in (i) on empirical grounds. When we ordered the ACWs of source-reconstructed resting-state time series of ROIs from low to high reflecting an ACW-based hierarchy, we observed that the TE values during both task states followed this hierarchical ordering. In accordance with our modeling results, total TE values of ROIs correlate negatively with the ACW values in the two MEG tasks. Together, our results demonstrate a direct relationship between the brain’s hierarchy of timescales and their inter-regional information transfer while being closely coupled together along a macroscopic uni-transmodal gradient in the cortex. This does not only complement and extend previous assumptions of the population-based constitution of the macroscopic timescale gradient^6^ but also carries major implications for cognitive processes^14^ and psychopathologies like schizophrenia^12,41^.

## Methods

### Computational model

We used a large-scale biophysical model that was introduced in^21^. A hierarchy of intrinsic neuronal timescales emerge from this model due to its architecture.

The model consists of the following system of differential equations:

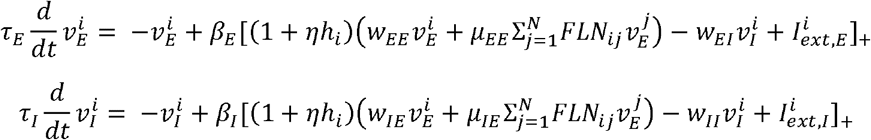

where *ν* is the firing rate, τ is intrinsic time constant, *I*_*ext*_ is the external input to the system governed by the slope *β* of the linear threshold f-I curve. *w* values are coupling parameters. *μ* is a fixed parameter that controls the strength of long-range excitatory input. (Fraction of Labeled Neurons) is the structural connectivity matrix based on a macaque study^22^ (supplementary figure 1). *E* and *I* correspond to excitatory and inhibitory, respectively; *i* and *j* denotes different regions. *h* values were assigned to each area such that the difference in values predicts FLN according to a logistic regression function *g*^−1^

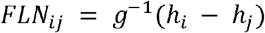

Resulting *h* values were normalized by dividing them to the largest *h*. Model parameters and various manipulations we made on the model were shown in table 1. We tried to explore a wide range of parameters while keeping the model’s outcome in a plausible range (i.e. not going to infinity or zero). The so-called default values were taken from Chaudhuri et al., 2015. The model was simulated for 300 seconds using Euler-Maruyama method with a time-step of 1 ms. The first 50 seconds were discarded. All simulations were performed 30 times. For different parameters, we used the same noises by resetting the random number generator.

**Table 1.** Model parameters and the changed values of them.

| Parameter | Default value | Manipulated values |
| --- | --- | --- |
| $w_{EE}$ | 24.3 pA / Hz | 12.15, 14.58, 17.01, 19.44, 21.87 |
| $w_{IE}$ | 12.2 pA / Hz | 13.42, 14.64, 15.86, 17.08, 18.3 |
| $w_{EI}$ | 19.7 pA / Hz | 21.67, 23.64 |
| $w_{II}$ | 12.5 pA / Hz | 10, 11.25 |
| $\beta_E$ | 0.066 Hz / pA | 0.033, 0.0396, 0.0462, 0.0528, 0.0594 |
| $\beta_I$ | 0.351 Hz / pA | 0.4212, 0.4914, 0.5616, 0.6318, 0.702 |
| $\mu_{EE}$ | 33.7 pA / Hz | 30.33, 32.015, 34.037 |
| $\mu_{IE}$ | 25.3 pA / Hz | 25.047, 26.565, 27.83, 30.36 |
| $\tau_E$ | 20 ms | 12, 16, 24, 28, 32, 36, 40 |
| $\tau_I$ | 10 ms | 6, 8, 12, 14, 16, 18, 20 |
| $\eta$ | 0.68 | 0.544, 0.5576, 0.5712, 0.5848, 0.5984, 0.612, 0.6256, 0.6392, 0.6528, 0.6664 |

Two different kinds of input were given to the model. First, we gave the sensory region V1 gaussian noise with 0-mean and unit variance. The other 28 regions received gaussian noise with a standard deviation of 10^−5^ and 0-mean representing their intrinsic dynamics. The second input scenario was similar to the task paradigm of empirical data and to also show the independence of the intrinsic uni-transmodal information transfer from the temporal structure of the extrinsic input. Starting from second 10, boxcars were created that lasted 2 seconds and had amplitude of 0.5. The duration between boxcars were 1 second. This resulted in 83 boxcars that spanned across simulation. These boxcars were added on the white noise and given to V1. We created trials by averaging the activity for the duration of the boxcar, and averaged the trials in order to obtain ERP-like data. Transfer entropy calculations were done on these ERP-like chunks.

### Estimation of Intrinsic Neuronal Timescales

We used the autocorrelation function to estimate intrinsic neuronal timescales (INTs) at resting condition. The autocorrelation function (ACF) *r* of a signal *x* is the signal’s correlation with itself on different time lags:

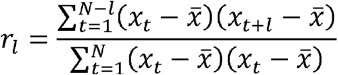

where *l* denotes lag, 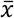 denotes the mean of *x* and *N* is the number of sampling points. We used ACW-0^14–16^ (the first lag below *r=*0) to extract one feature per signal that is representative of INT. We picked ACW-0 instead of ACW-50^10,14,17^ (the first lag below *r* = 0.5) because it was shown to better differentiate different regions in the brain^16^

### Transfer Entropy

Transfer entropy (TE) is a non-parametric model-free measure of information transfer, that is, effective connectivity^25,27^ that is shown to be informative in neural data^28–31^ as well as other fields^42– 44^. For two time series *x* and *y*, transfer entropy from *y* to *x* is the Kullback-Leibler divergence from the generalized Markov property 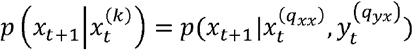. Here q_xx_ and q_yx_ refer to the Markov’s lags: 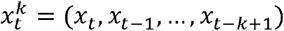 and *p* denotes probablity. The general formula for. transfer entropy is given by

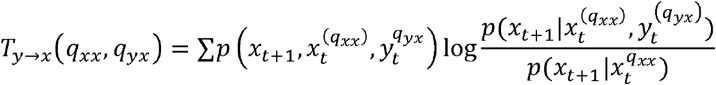

Where *T* _*y→x*_ quantifies information transfer from *y* and *x*. In other words, how much our uncertainity about the next state of *x* reduces when we know the previous *q*_*yx*_ values of and previous *q*_*xx*_ values of compared to only knowing previous *q*_*xx*_ values of .*x* TE values were calculated with the RtransferEntropy package^27^.

Note that this calculation assumes discrete data. In order to discretize our continuous time series, we used symbolic encoding method. We used the default methods in RtransferEntropy package for the choice of parameters^27^ which discretizes the data into 3 bins using 5% and 95% empirical quantiles of the data, resulting in three symbols in the end where first (third) symbol represents lowest (highest) values. Other discretizations (bins using 20, 40, 60 and 80th percentiles as well as 25th, 50th and 75th percentiles) gave qualitatively equivalent results (not shown).

The rationale for using TE as opposed to other metrics of effective connectivity is twofold. As opposed to methods based on dynamic causal modelling^45,46^, transfer entropy is model-free. No a priori model selection is required, making it more flexible. As opposed to Granger causality^47,48^, nonparametric metric TE can capture nonlinear relationships which Granger causality underestimates^27^.

### Data-Driven Determination of Markov Order

One challenge in using TE is the determination of Markov lags. Usually, experimenters set one Markov lag to a fixed value (either one or equal to the other Markov lag) and try to determine others using false neighbourhood approaches^49,50^. Other methods include using certain instantaneous phase properties of the signal and fixing the Markov lag same for both variables^51^, and fixing the first Markov lag to the *q*_y*x*_ that minimizes the mutual information between *x*_*t*_ and 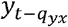 while setting the other Markov lag *q*_*xx*_ to 1^52,53^. While these methods are computationally efficient, they might over or under estimate the cases where Markov lags are different between driving and driven systems. In order to determine Markov orders, we used a data-driven method^54^. This method builds multivariate autoregressive (MVAR) models with different Markov lags and compares them using generalized Bayesian Information Criterion (gBIC) to pick the model that has the least complexity and explains highest variance. Then, the estimated Markov lags are extracted from the best MVAR model to be used in TE estimation.

A MVAR model of the form

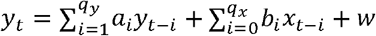

Where *w* denotes the error term. This can be represented with linear algebra notation as

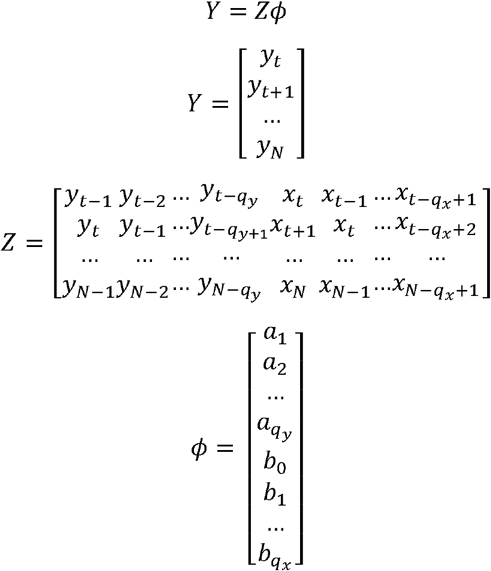

where *t* = max(*q*_*x*,_*q*_*y*_*) +*1 and *N* denotes end of the array. Since *Y* and *Z* is known from the data, *ϕ* can be estimated using ordinary least squares method:

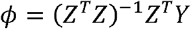

We can generalize the MVAR model to two equations that influence each other:

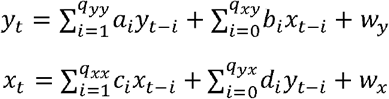

and solve *ϕ* for *y* and *x* seperately using the method above. The covariance matrix of the solution is given by:

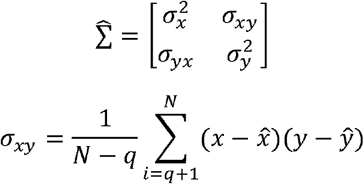

Where 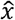 and ŷ are estimated values according to the estimated AR coefficients. Finally, gBIC value is computed as 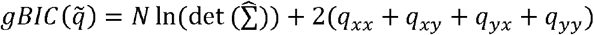 In *N* where 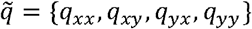 The 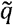 set that minimizes gBIC was chosen for TE calculation. gBIC was chosen instead of BIC since it doesn’t rely on strict criteria^55–57^.

We tried values from 1 to 20 for every value in set .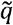 To achieve this, every computation has to be performed 20^4^ times if one uses a brute-force method. To decrease computation time, we used the so-called “greedy” strategy described in^54^

1. Use gBIC to obtain the maximum order *q*_*max*_ and select this order for the range for further calculations. Here, assume that *q*_*xx*_= *q*_*xy*_ = *q*_*yx*_ = *q*_*yy*_ = *q* and try *q* values from 1 to 20 to get the *q*_*max*._
2. Evaluate *q*_*i,j*_ of *X* while setting *q*_*ij*_ for *Y* fixed to *q*_*max*_ determined in step 1. For both lags for *X*, evaluate orders from 1 to *q*_*max*_ in a double for-loop and pick the ones that minimize gBIC.
3. Use the *q*_*ij*_ for *X* obtained in step 2 while evaluating *q*_*ij*_ from 1 to *q*_*max*_ determined in step 1 akin to step 2 to get the final *q*_*ij*_ values.

It might be argued that this method of determination might bias TE values. To counter this argument, we correlated total TE values of different ROIs with their respective between (*q*_*xy*_+ *q*_*yx*_) and within (*q*_*xy*_+ *q*_*yy*_) total Markov lags for the model and two MEG tasks. The results can be in supplementary figure 2.

**Figure 2.**
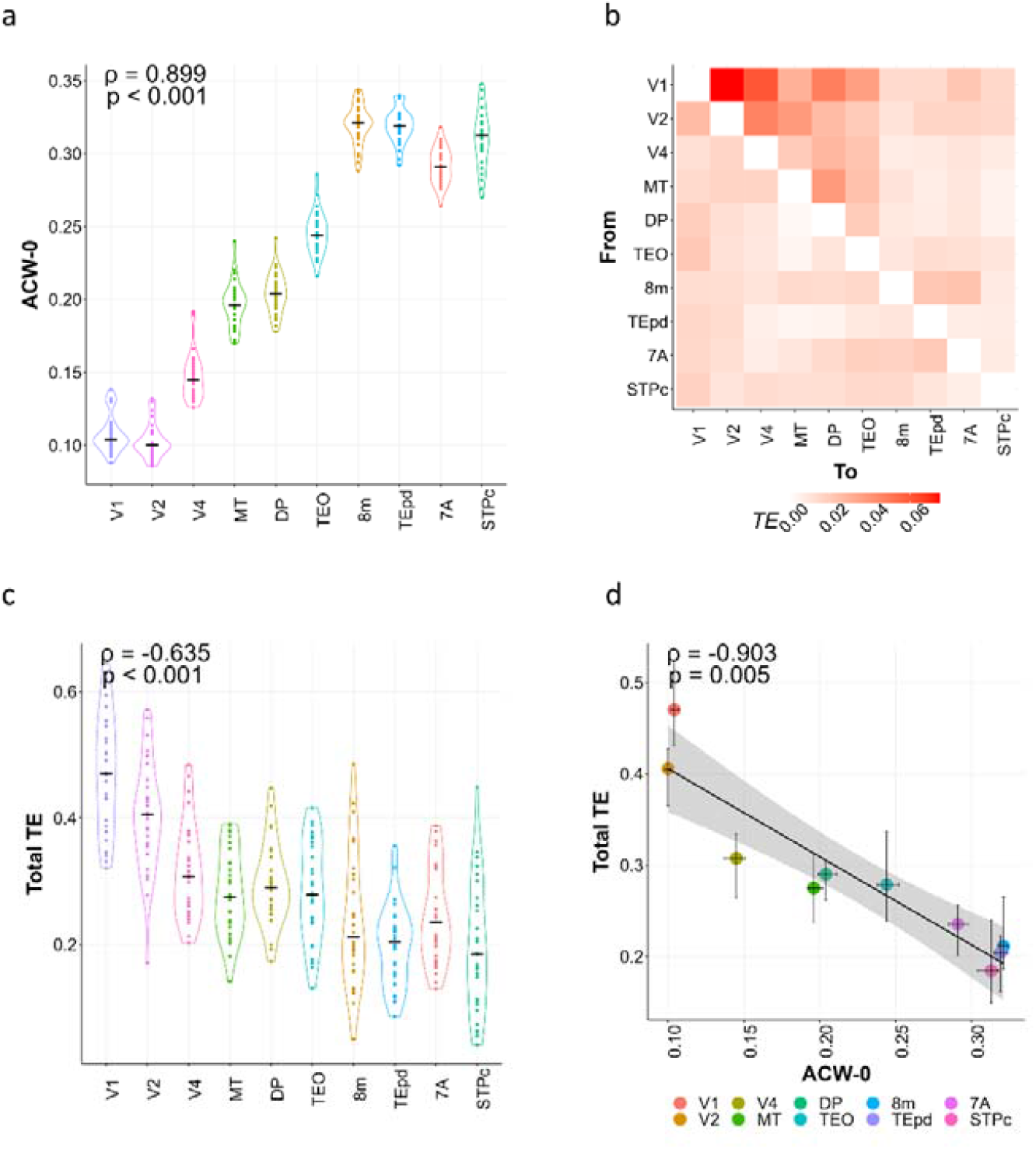
Results from the model with parameters described in Chaudhuri et al., 2015. A. A hierarchy of intrinsic neural timescales (INT) with shorter INT at early sensory areas and longer ones at transmodal areas. Correlation results rho and p show the Spearman correlation between ROI orderings and their respective ACW-values. B. Matrix of transfer entropy (TE). C. Total TE of the regions follow the hierarchy of timescales with higher TE at lower areas and vice versa. D. ACW-0 values negatively correlate with TE values. Dots show median, 95% percentiles are estimated using bootstrapping. n = 30 simulations for every manipulation in each panel. The correlations were done on medians of 30 simulations.

### MEG Data Acquisition and Experimental Details

We used publicly shared resting-state and working memory task magnetoencephalogram (MEG) data from Human Connectome Project (HCP)^33–35^. The details about acquisition parameters can be found in^35^. The data had 89 age age and sex matched subjects for the resting state scan, 83 for the working memory task and 61 for motor task. For resting state, subjects were instructed to lie quietly with eyes fixed on a cross. Three successive resting state runs (5 minutes each) were collected. The 2-scan working memory task is an alternating 0-back and 2-back task in which subjects were presented with pictures of tools and faces. The stimuli were presented for 2 seconds following a 500 ms fixation. In the motor task, participants were asked to move their left or right hands or foot depending on the visual stimuli presented. The stimulus was presented for 150 milliseconds, interspersed with 1050 milliseconds where the subject can move their limbs. The exact details about the task designs can be found in^35^.

### MEG Preprocessing

We used the preprocessed data from HCP at *rmegpreproc* and *tmegpreproc* stage of their pipeline ^35^ for rest and task respectively. For *rmegpreproc*, briefly, channels with low correlations with their neighbors and a high variance ratio were removed; artefactual components and components that are results of non-neuronal processes (such as eye blinks, muscle or sensor artifacts) were removed based on an automated iterative ICA procedure. Different from HCP procedure, bad segments that were identified on z-scored amplitude, flatness (lack of signal) and muscle artifacts were linearly interpolated instead of removing in order to maintain signal’s temporal continuity^58^. *tmegpreproc*, in addition to the steps of *rmegpreproc*, divides the data into trials. Here, since we were interested in an event-related analysis, we rejected the trials with bad data.

To do source reconstruction, we used the *icamne* step of HCP processing pipeline for both rest and task using the independent components identified at *icaclass* step for both rest and task. In *icaclass*, independent component decomposition is repeated several times with different initial conditions and the decomposition with the lowest artifact residual and largest number of brain components is retained as the best decomposition. The *icamne* applies weighted minimum norm estimation to project sensor maps of the brain independent components into the source space which consists of individual surface-registered cortical sheets of 8004 vertices.

As a final step, we spatially interpolated resulting data to match the HCP-MMP parcellation atlas^59^ and used ft_virtualchannel function from fieldtrip to obtain one time-series per region using the singular value decomposition method.

We averaged the trials in the task data across two scans to get event-related fields. TE values were calculated on ERFs. For resting state ACW calculations, we used a method similar to^17^. We cut the resting state data to non-overlapping 2 second segments (matched to task ERP duration) in 3 resting state scans, calculated one ACF per segment, averaged those ACFs and extracted ACW-0 from the averaged ACF. Thus, we obtained one ACW-0 value per region per scan per subject. We averaged ACW-0 values across scans to get one ACW-0 per region per subject.

### Selection of Regions of Interest – Model

In the model, we selected regions that form the visual hierarchy based on^60^ since it used the same anatomical data we used in the model. We ended up with using regions V1, V2, V4, TEO, Tepd, MT, DP, 8m, STPC and 7A.

### Selection of Regions of Interest – MEG

We determined the regions of interest for the MEG working memory task based on ^36^. This paper used working memory task from HCP functional magnetic resonance imaging (fMRI) data with the same task design and temporal characteristics as the MEG one we used. Among 160 regions from an atlas^61^ and 4 added subcortical regions, the authors calculated psychophysiological interactions (PPIs) for each pair of ROIs, then treated those PPI matrices as network graphs where each ROI represents one node and PPI effects represent edges of the graph. They calculated node degree as number of significant edges per ROI. We took 9 regions that had the highest node degree and found the corresponding regions from the Glasser atlas based on their MNI coordinates. Thus, we ended up with the following regions: R_7pM, R_LIPv, R_V3B, R_7PC, L_TPOJ3, R_6d, R_OP1, L_46 and L_V3.

The regions for the motor task were determined based on the work of Saetia et al^37^ which used the motor task from HCP fMRI data, again with the same task design and temporal characteristics as the MEG task. The authors performed univariate group subtraction analysis to find significant clusters of activity. We used the table 2 in their paper and found equivalent regions in the Glasser atlas to be used in our study. Final 12 regions were L_4, L_6v, R_4, R_6v, L_6d, L_1, R_6d, R_V1, L_6mp, L_V2, R_6mp, R_V2.

## Results

### Modelling – A Hierarchy of Information transfer follows the Hierarchy of Intrinsic Neuronal Timescales

We started our analyses by replicating the model in^21^. Similar to resting state and task paradigms in human neuroimaging, we first stimulated V1 with an excitatory gaussian noise as an input and gave all other regions relatively small excitatory gaussian noise. This represented the resting state and we calculated ACW-0 values in this state. For task, we added boxcar inputs on top of the same noise and averaged the durations of neural activity during these boxcars to obtain an ERP-like structure and calculated TE on this state. As in the results of Chaudhuri et al^21^, a hierarchy of timescales emerged from an excitatory stimulation of V1 with gaussian noise. We found lower ACW-0 at early sensory areas and higher ACW-0 in multimodal areas, (figure 2A) (spearman correlation between ROI orderings and ACW: *ρ*= 0.899, p < 0.001).

We next calculated the information transfer of time series from every region pair using a nonparametric model-free measure of effective connectivity called transfer entropy (TE) while stimulating V1 with gaussian noise and adding boxcars on top of that, subsequently averaging activity in those boxcars akin to an event-related analysis. In our TE calculations, we used a data-driven method to determine Markov lags. To make sure that our method does not bias estimated TE values, we correlated estimated total Markov lags with total TE values for the model and 2 MEG tasks. None of the correlations were significant (details can be seen in supplementary figure 2). This shows that regardless of the model or data, our TE calculations are not driven by Markov lag estimation method. The resulting directional matrix of TE is given in figure 2B. One striking feature of this matrix is the higher information transfer in early sensory areas versus lower information transfer in higher associative areas. To quantify this feature, we summed all the incoming and outgoing transfer entropy for every region. The resulting plot is given in figure 2C. As can be seen from the figure, there seems to be a hierarchy of the information transfer similar to the INT hierarchy (spearman correlation between ROI orderings and ACW: *ρ*= -0.635, p < 0.001).

To investigate how the INTs relate to the observed directional information transfer, we plotted the median ACW-0 values of all the regions against the median of their total TE values (sum of incoming and outgoing TE) (figure 2D). The result showed a strong negative correlation (*ρ* = -0.903, p = 0.005): the longer the ACW indexing longer temporal window, the lower the TE indexing low information transfer. This indicates that the hierarchy of information transfer follows the hierarchy of intrinsic neuronal timescales in a negative way.

Together, these results indicate that INTs and information transfer are negatively related. Does the topographic hierarchy of INT causally relate to the information transfer? To address this question, we next manipulated various model parameters of the model in order to study their impact on the INT hierarchy and its relationship with the information transfer.

### The Hierarchy of Information transfer Depends on the Hierarchy of Timescales I – Changes in Excitatory and Inhibitory Coupling Parameters

In order to probe mechanistic interpretations of the INT-mediated hierarchy of information transfer, we manipulated different parameters of the model. We started with local excitatory-to-excitatory connection strength. We defined the existence of a timescale hierarchy as a positive and significant Spearman correlation between ROI orderings and ACW-0 values (i.e. 1, 2, 3, …; ACW-0 value of V1, V2, V4, …). A significant and positive correlation means increasing ACW-0 values along the ROIs. The results of the correlation tests are detailed in figure 3A. Simulations showed that even a small reduction in the parameter is enough to break the timescale hierarchy (figure 3A, = 21.87); as only the default value in the original model showed the timescale hierarchy according to the definition we sketched above. If we define the hierarchy of information transfer in the same way, even in the loss of hierarchy of timescales, the hierarchy of information transfer seems to be intact even though it is not consistent (i.e. doesn’t continuously decrease) (see figure 3B for correlation results).

**Figure 3.**
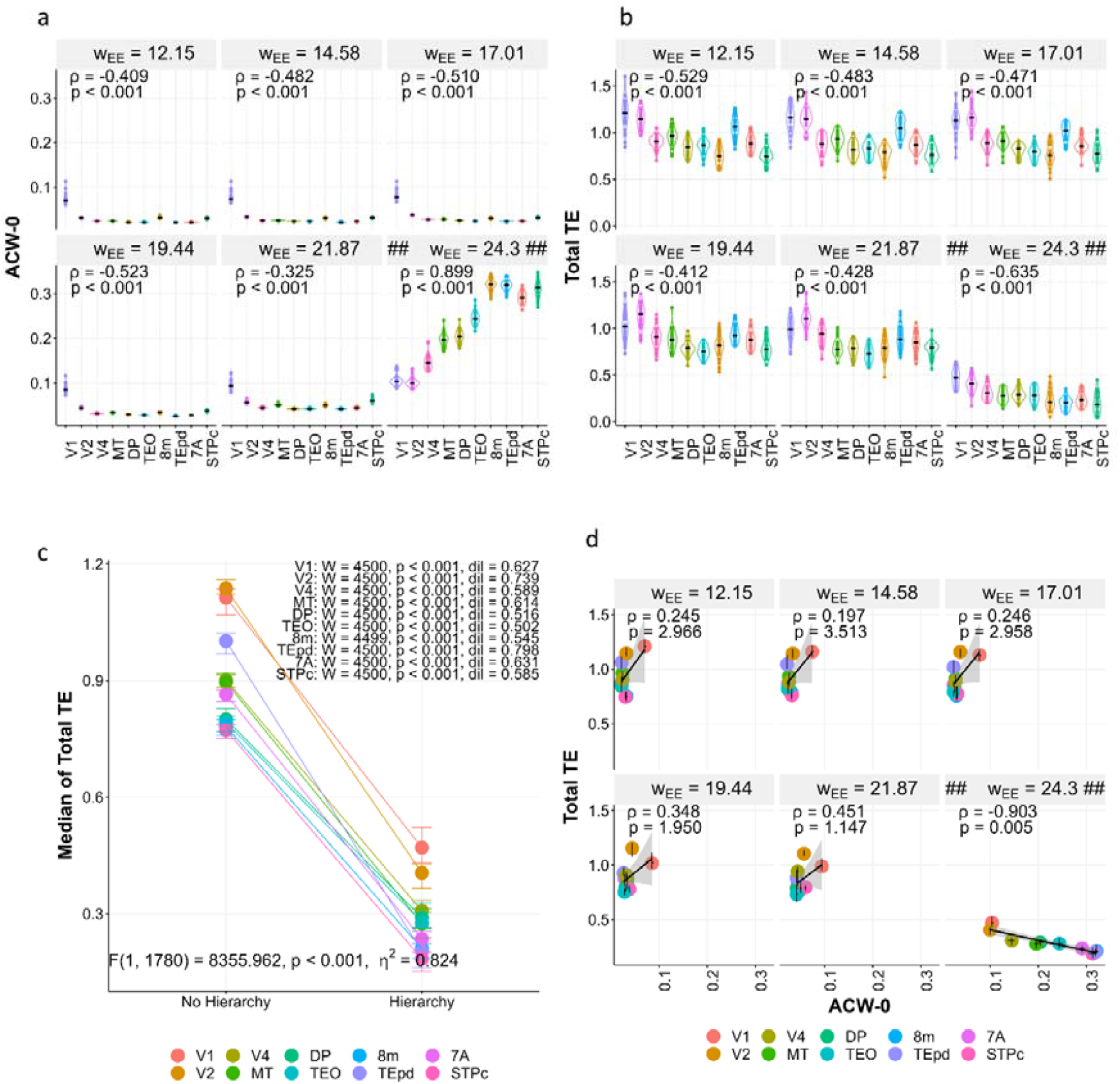
Manipulation of local excitatory-to-excitatory coupling parameter. A. ACW-0 values for changing. Correlations are the same as figure 2. B. Total TE (summation of incoming and outgoing) values for changing. Correlations are the same as figure 2. C. Total TE increases when the hierarchy is lost. We defined the existence of hierarchy as a positive and significant correlation between ROI orderings and ACW-0 values. W: Wilcoxon test statistic. dil: effect size measure difference in location. F value and corresponding statistics p and partial are results from a two-way ANOVA with variables ROIs and the existence of hierarchy. Values show the results for the dummy variable existence of hierarchy. D. Correlations between ACW-0 and Total TE. In all figures, the panels with ## = 24.3 ## shows the value in figure 2. n = 30 simulations for every manipulation in each panel. The correlations were done on medians of 30 simulations.

To compare total TE in networks where hierarchy was lost versus preserved, we grouped different manipulations into two groups: hierarchical vs. non-hierarchical (see the first paragraph of this heading for criterion.). The networks where hierarchy was lost had higher total TE compared to the networks where hierarchy was preserved (2-way ANOVA, F(1, 1780) = 8355.962, p < 0.001, partial *η*^2^ =0.824; post-hoc comparisons can be seen in figure 3D). The overall TE matrix (similar to figure 2B) is given in supplementary figure 10a.

The *w*_*EE*_. based disruption of the INT hierarchy also affected the relationship between information transfer and INTs. When the timescale hierarchy was lost, the correlation between ACW-0 and total TE was also lost (figure 3D). The only significant relationship was on the default value of the parameter.

Finally, analogous results were obtained for other related local inhibitory and excitatory parameters in the model like *w*_*EI*_, *w*_*EI*_ and *w*_*II*_ (supplementary figures 3, 4, 5 respectively). Importantly, we again saw that minimal changes in all three model parameters lead to the disruption of both INT hierarchy (supplementary figures 3, 4, 5 respectively). Importantly, we again and the corresponding hierarchical regional differentiation of the information transfer (TE). This again was accompanied by the disruption of the relationship of ACW and TE.

### The Hierarchy of Information Transfer Depends on the Hierarchy of Timescales II – Changes in Other Model Parameters

We next asked ourselves whether the hierarchical information transfer may be related to other model parameters that affect both local and global connections: intrinsic time constants for excitatory and inhibitory populations *τ*, slopes of the rectifying curves *β*, and the constant that scales the hierarchy of excitatory connections *η*.

We probed *β*_*E*_, the slope of the rectifier curve for excitatory population. For a small change in *β*_*E*_ (from 0.06 to 0.054), the hierarchy of timescales was preserved but was slightly lower (figure 5A, two plots in the lower right corner). However, in any further reduction of *β*_*E*_, it was lost (figure 5A, all the other plots). The resulting TE matrices in different model manipulations are given in supplementary figure 10e. Similar with the trend above, the existence of a TE hierarchy seems to be independent of ACW hierarchy (figure 5B) while the relationship between ACW and total TE (across all regions of the hierarchy) depends on the existence of a timescale hierarchy (figure 5D). As above, a loss of hierarchy brought an increase in total TE in all ROIs (F(1, 1780) = 277.44, p < 0.001, partial *η*^2^ =0.135, figure 5C). The equivalent results for *β*_*I*_ are described in supplementary material (supplementary figure 7).

We also changed the time constants of excitatory and inhibitory populations and. The timescale hierarchy was fairly resistant against manipulations in, no manipulation in our range was able to fully disrupt it even though at = 12 and 16, the timescales of intermediate regions are flat (supplementary figure 8A). Finally, in all the different manipulations, the negative correlation between ACW and total TE was preserved; in accordance with the general pattern (supplementary figure 8C). Similar results for can be seen in supplementary figure 9. The TE matrices for and are shown in supplementary figure 11a and b, respectively.

Finally, we changed the parameter that controls the gradient of excitatory connections across regions. The rho value of the spearman correlation between ROI orderings and both ACW / TE increases / decreases with increasing (figure 6A and B, supplementary figure 12 shows the scatterplot between and rho values.). The TE matrices of manipulations are shown in supplementary figure 11c. The total TE values of ROIs and their correlation to ROI orderings are given in figure 6B. Interestingly, in some of the manipulations, even though ACW hierarchy was preserved, the correlation between ACW and TE became non-significant (figure 6C). However, we should note that rho values in spearman correlations were very high (∼-0.65) even in the nonsignificant results. This might be the result of using very strict Bonferroni correction.

In our simulations, we saw that value can control the steepness of the timescale hierarchy. Thus, we wanted to test the relationship between steepness of the TE and ACW hierarchy using the different manipulations in. We plotted both steepness values calculated as the slope of the linear fit between ROI orderings and ACW/TE values, (figure 6D). The spearman correlation between two variables was negative and significant (= -0.945, p < 0.001). This shows that the steeper the timescale hierarchy, the steeper the TE hierarchy.

In order to probe further our hypothesis about the dependence of the total TE on the timescale hierarchy, we summed all TE values across ROIs for every manipulation we made in the model and correlated it with the corresponding slopes of the ACW hierarchy obtained via a linear regression between ROI orderings and ACW values (figure 7A). The correlation was negative and highly significant (r = -0.76, p < 0.001): when the timescale hierarchy gets steeper, total information transfer decreases – that further supports the close relationship of INT hierarchy and information transfer.

In figure 7A, we saw that many values are at the left and right edges of the plot, at the extremes of ACW slopes. To look into this closer, we plotted the histograms of ACW slopes (figure 7B) and total TE values (7C). The ACW slopes show a more bimodal distribution, indicating a jump from the hierarchical state to non-hierarchical state (7B). The total TE on the other hand seems to have a more uniform distribution (7C). This echoes the decoupling of TE and ACW hierarchies that we saw in the other figures (for example, figure 2A and B).

Taken together, our findings show that changes in the model parameters that are not related to excitatory and inhibitory coupling (and)disrupt the INT hierarchy and its negative relationship with the information transfer. While the findings resemble those of the changes in the excitatory – inhibitory coupling parameters, there was slightly more ‘tolerance’ for the changes in these parameters, preserving the INT hierarchy and its relationship with the information transfer in the sense that one needs larger changes in those parameters to break the timescale hierarchy. Together with the excitatory-inhibitory findings, these observations clearly support the assumed causal role of INT hierarchy in differentiating the information transfer across the different regions.

### Empirical data – information transfer follows the hierarchy of timescales

Using modelling, we so far demonstrated that the information transfer is negatively related with the INTs .. Does this also hold in empirical data? Unlike the computational modelling, we did not assume an a priori known topographic hierarchy of uni- and transmodal regions. Instead, using the HCP MEG dataset^33–35^, we determined the relevant regions by their known degrees of recruitment^36,37^ during two tasks, e.g., working memory and motor task. This enabled us to investigate that even in the absence of an a priori defined topographic hierarchy (as in our model), there is a negative relationship of INT (ACW) and information transfer (TE) for the task-relevant regions. Similar to the model, we calculated ACW-0 values at the resting state and TEs at task.

For the working memory task, we ordered the regions according to their ACW-0 values at rest (figure 8A). As expected, this showed a significant and positive Spearman correlation (*ρ* = 0.457, p < 0.001) between the ROI orderings and their ACW values. The resulting TE matrix can be seen in figure 8B. The total transfer entropy values for the regions followed the ACW hierarchy and were consistent with the simulation results. While it was not as pronounced as in the simulations, we saw a significant negative correlation between the ACW-based ROI orderings and median total TE values (rho = -0.179, p < 0.001) (figure 8C). To show that our way of estimating ROI orderings does not give random results, we randomized the order of ROIs 10000 times and fit linear regression curves between ROI orderings and median total TE values. This created null distributions for us. Our ACW-0 based linear regression slope had a 2-sided p-value (calculated as the probability of getting as extreme or more extreme slope in the null distribution of randomized ROI orderings) was 0.0074 (the histogram can be seen in supplementary figure 13, see also supplementary figure 13 for the same analysis done for motor task). Similar to our models, median of total TE values per ROI correlated negatively with their median ACWs: the longer the ACW of a region, the lower its total TE, e.g., its information transfer (rho = -0.75, p = 0.025, figure 8D). Similar results for the motor task can be found in supplementary figure 14. These results strengthen the relationship we observed on the model regarding the INT hierarchy and its close relationship with the information transfer.

## Discussion

In this analysis, we probed the relationship between information transfer and the hierarchical nature of intrinsic neural timescales (INTs) using both modeling and empirical data. Our results can be summarized in three points: (i) total information transfer follows the hierarchy of INTs, (ii) total information transfer increases when the hierarchy of INTs is lost as based on the changes in excitatory and/or inhibitory model parameters, (iii) total information transfer is negatively correlated with the steepness of the hierarchy.

### Hierarchy of information transfer follows the hierarchy of intrinsic Neural Timescales

Using the information-theoretical measure transfer entropy^25,27^ (TE), we quantified information transfer between our regions of interest. TE is a nonparametric and model-free measure of effective connectivity, thus, it can capture nonlinear relationships between time series without assuming an a priori model^27^. We used a data-driven method to determine the lags in the calculation of TE independently for all variables^54^. This way, we could capture the heterogeneity in time dependencies within and between regions; a point that we deem important in a study that is aiming to relate TE to temporal correlations, e.g., ACW as a proxy of INT.

We found that just like ACW, TE also follows a sensory-to-associative hierarchy with higher TE in lower-order areas and vice versa. This adds to what we know about the macroscopic gradients along the rostro-caudal axis of the brain^6^. Previous studies found that a) synaptic excitation increases^4^; b) inhibitory gating of input increases^5^; and c) INTs increase^3,7,8^ from sensory periphery to associative core of the brain.

Following these observations, we hypothesize that our findings are an effect of inhibition and INTs working against long-range synaptic excitation: Increasing inhibitory gating along the gradient reduces the information transfer even though the synaptic excitation increases over the hierarchy. This is achieved by local inhibition’s counter-balancing effect on long-range excitation.

One example of this effect is given in the case detailed in figure 4E and 4F. Here, we have a decrease in global excitation but this results in an increase in information transfer. One possible explanation is that in the case with reduced global excitation and a lack of hierarchy, inhibitory activity cannot compensate for excitation since global excitation also excites inhibitory populations indirectly. We see this in figure 4F.

**Figure 4.**
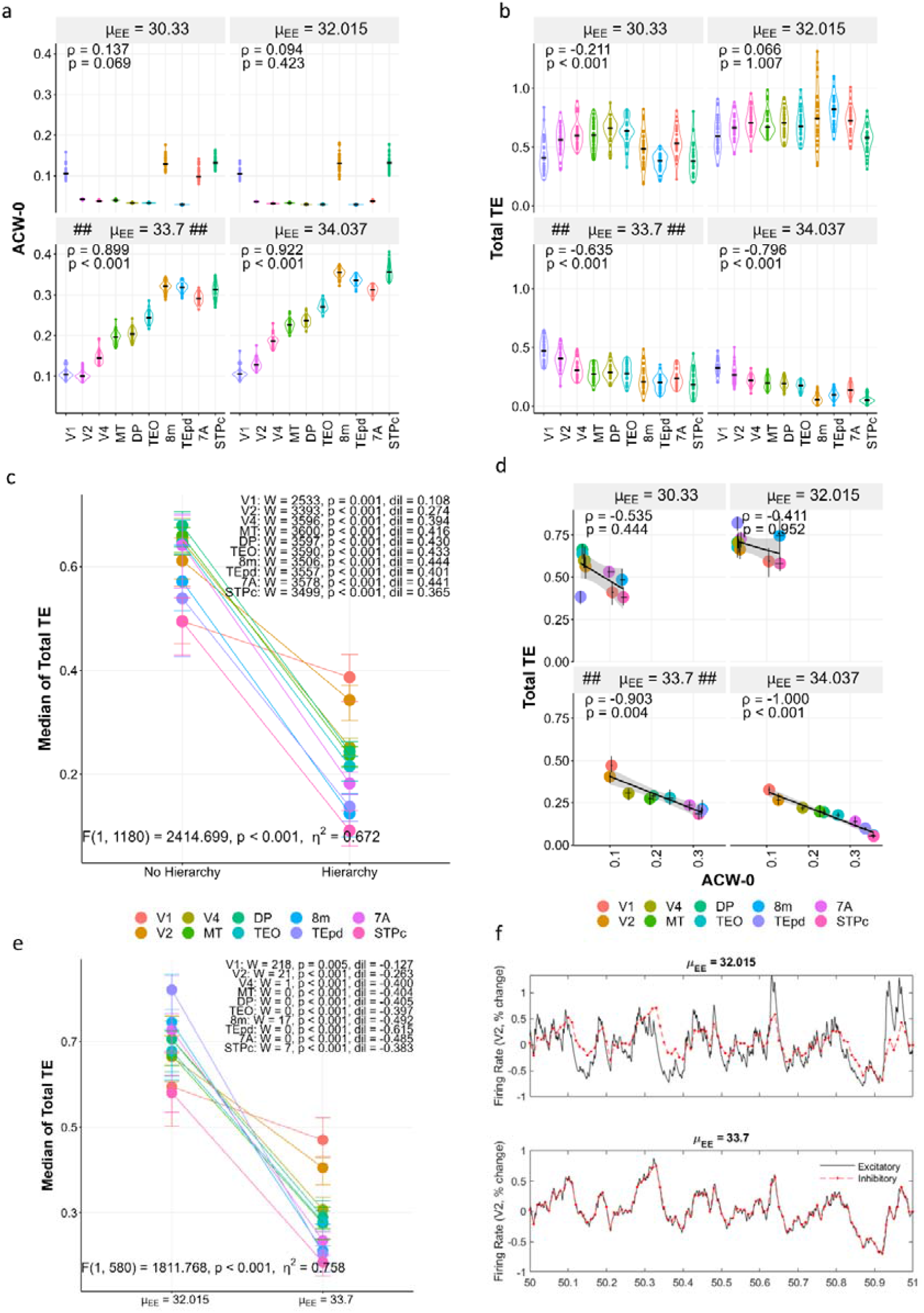
Manipulation of global excitatory-to-excitatory coupling parameter μ_EE_. A. ACW-0 values for changing μ_EE_. Correlations are the same as figure 2. B. Total TE (summation of incoming and outgoing) values for changing μ_EE_. Correlations are the same as figure 2. C. Total TE increases when the hierarchy is lost. We defined the existence of hierarchy as a positive and significant correlation between ROI orderings and ACW-0 values. W: Wilcoxon test statistic. dil: effect size measure difference in location. F value and corresponding statistics p and partial η ^2^ are results from a two-way ANOVA with variables ROIs and the existence of hierarchy. Values show the results for the dummy variable existence of hierarchy. D. Correlations between ACW-0 and Total TE.. E. A decrease in μ_EE_ causes an increase in total TE. F. Time series of excitatory and inhibitory firing rates in one example region V2 for the two μ_EE_ values in E, reflecting percent change from mean. n = 30 simulations for every manipulation in each panel. The correlations were done on medians of 30 simulations.

We next continued with the manipulation of the global excitatory-to-excitatory coupling strength. For a slight increase of (1%), the timescale and information transfer hierarchy were preserved whereas for more severe reductions, it was lost (figure 4A and B). Supplementary figure 10 shows the TE matrix in different manipulations. Similar to results, the networks where hierarchy is preserved has lower total TE in all regions compared to the networks where hierarchy was lost (figure 4C). In the manipulations where timescale hierarchy was preserved, we saw the negative correlation between ACW and total TE whereas in all the manipulations where it was lost, their correlation was also lost (correlation results are given in figure 4D). The results for can be seen in supplementary material and in accordance with the aforementioned results (supplementary figure 6).

Interestingly, decreasing global connectivity strength did not cause a decrease in information transfer, on the contrary, it caused an increase. We probed this via a 2-way ANOVA (F(1, 580) = 1811.768, p < 0.001, partial *η*^2^ = 0.758) and subsequent post-hoc comparisons (figure 4E). In every region, the median of total TE increased significantly for a decrease of *μ*_*EE*_ from 33.7 to 32.015. One instance of the time series of excitatory and inhibitory populations in V2 is given in figure 4F where we see that in the network with lower global coupling, local inhibition can’t catch up with excitation. This is in accordance with the predictions of Li and Wang^32^.

The mechanism of INTs on information transfer is not so straightforward. Echoing a previous review by Golesorkhi et al^2^, (see figure 7 in that paper) INT of a brain region can serve as the “sampling rate” in digital signal processing terms: the rate in which one can capture information where each sample is one time-point. The sampling rate determines the highest frequency (Nyquist) information that can be captured by the recording device^40^ with a relationship Nyquist = sampling rate / 2. Any information that has a higher frequency than the Nyquist will not be able to be captured by the recorder. In our case, different regions in our model have to encode the information coming from other regions. At every step along the hierarchy, high frequency information of the lower-order regions will be gradually lost along the uni-transmodal hierarchy, resulting in downsampling and decrease in information transfer of the higher-order regions.

Given that this is exactly what we observed in both computational and empirical data, our results showing attenuation of information transfer along the timescale hiearchy are well compatible with such down-sampling as supposedly underlying mechanism. This is further supported by the fact that the elimination of the timescale hierarchy, through manipulations of the model parameters, lead to an overall increase in information transfer. Albeit tentatively, we therefore assume that downsampling may be a key mechanisms modulating the observed attenuation of information transfer along the uni-transmodal timescale hierarchy.

Going even one scale below, neurons can be thought as analog-to-digital converters. They accumulate continuous / analog information from environment and when a threshold is crossed, they fire a digital (0 or 1: not fire or fire) action potential: the basic unit of information in the nervous system. A long INT will cause the neuron to respond later to the environmental input and thus, making it miss the high-frequency information; effectively acting as the sampling rate. As a result, high frequency information will be gradually lost when it is propagated from lower-to-higher order areas, acting as a natural low-pass filter along the rostro-caudal gradient of the brain. Such an interpretation is supported by our particular example in figure 4E; we can hypothesize that the loss of timescale hierarchy enabled the overall increase in information transfer in the model even though the global excitatory-to-excitatory coupling strength is increased.

We captured the same relationship in human MEG data during both working memory and motor tasks as well. When the source-reconstructed MEG time series were ordered based on their resting state autocorrelation window (ACW) values, their TE during task followed the same order, and correlated with ACW negatively; confirming the theoretical predictions of the model in behaving humans. We hope to see upcoming studies that strengthen this link in empirical analyses.

### Total information transfer increases when the hierarchy of timescales is lost

Inspired by the theoretical work of Li and Wang^32^, we manipulated different parameters of the model and calculated ACW and TE again. In their paper, using perturbation theory, Li and Wang defined three necessary conditions for a hierarchy of timescales. The first condition is small 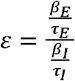, which corresponds to distinct electrophysiological properties of excitatory and inhibitory populations with larger f-I slope and smaller membrane time constant for inhibitory populations. The second one is small 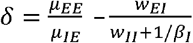.This ensures that any long-range excitation coming from other regions has to be countered by the local inhibitory population. The final condition is *h*: a gradient of increasing synaptic excitation that scales both local and long-range excitation.

Different manipulations we made on the model disrupt these conditions and break the timescale hierarchy. Importantly, regardless of the parameter that we changed (whether it is slope of the rectifier function, or local excitatory-to-excitatory coupling or any other one); when the timescale hierarchy was lost, this came with an overall increase of information transfer.

We should note that the timescale hierarchy was more resistant against disruptions in *ε* compared to disruptions in *δ*. For example, in supplementary figure 8, even when we decreased *τ*_*E*_ by 20% (*τ*_*E*_ = 16 which corresponds to a 25% increase in *ε*) the hierarchy of timescales persisted. On the other hand, even a 5% decrease in *μ*_*EE*_ (see figure 4 with *μ*_*EE*_ = 32.015; results in 12% increase in *δ*) was enough to break the timescale hierarchy. Although this has to be confirmed empirically, we hypothesize that local inhibition of long-range excitation is much more crucial in constituting the timescale hierarchy compared to electrophysiological properties of excitatory and inhibitory populations.

Another finding to note was, a 30% decrease of *β*_*E*_, (figure 5, *β*_*E*_= 0.042) was also enough to break the timescale hierarchy; which corresponds to a 30% decrease in *ε*, working against the small *ε* condition. We should note that, in a brilliant move, Li and Wang deliberately ignored the rectifier function in order to make the model analytically tractable since in the default parameters, the firing rate is always non-zero indicating that the rectifier function is not in effect. This reduced the model to a linear model which enabled them to solve the equations. In their work, Li and Wang claim that numerical results might contradict with their predictions due to this discrepancy. We hypothesize that this discrepancy between *ε* and *δ* might be due to the effect of rectifying curve in networks with no timescale hierarchy.

**Figure 5.**
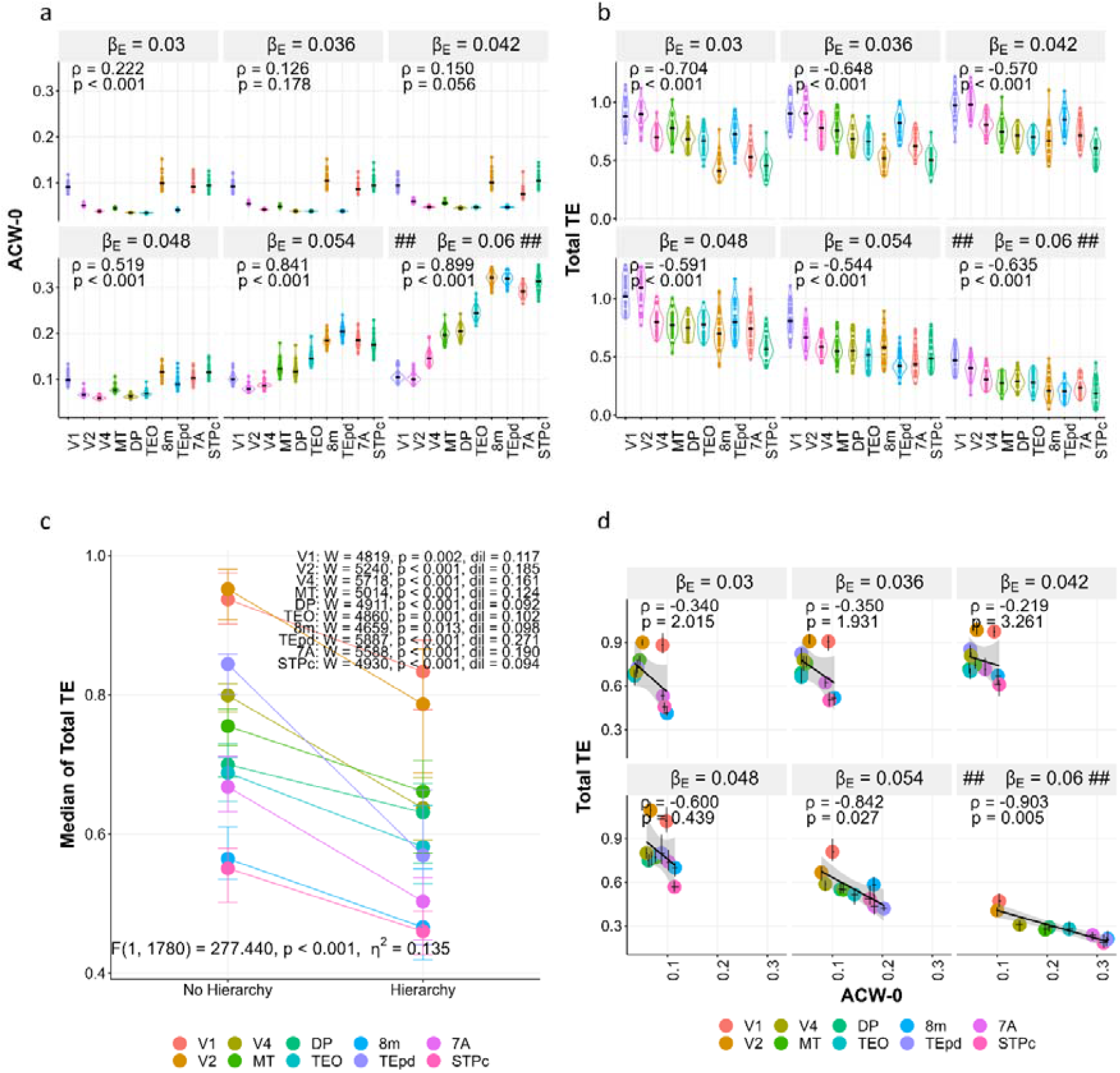
Manipulation of the slope of f-I curve. A. ACW-0 values for changing. Correlations are the same as figure 2. B. Total TE (summation of incoming and outgoing) values for changing. Correlations are the same as figure 2. C. Total TE increases when the hierarchy is lost. We defined the existence of hierarchy as a positive and significant correlation between ROI orderings and ACW-0 values. D. Correlations between ACW-0 and Total TE. W: Wilcoxon test statistic. dil: effect size measure difference in location. F value and corresponding statistics p and partial are results from a two-way ANOVA with variables ROIs and the existence of hierarchy. Values show the results for the dummy variable existence of hierarchy. In all figures, the panels with ## = 0.06 ## shows the value in figure 2. n = 30 simulations for every manipulation in each panel. The correlations were done on medians of 30 simulations.

**Figure 5.**
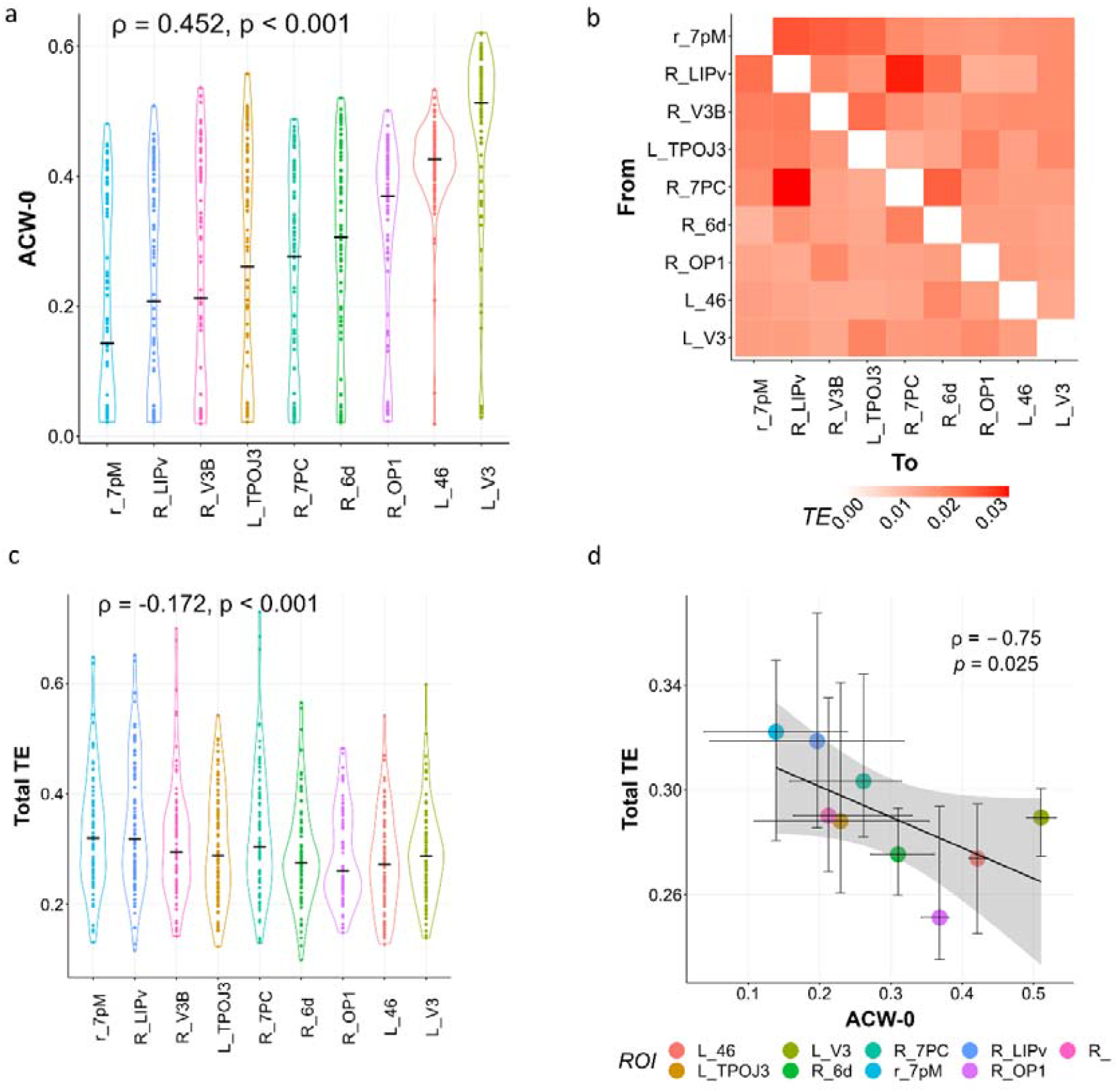
Replication of results in the HCP-MEG dataset working memory task. A. Resting state ACW-0 values, ordered from lowest to highest (n=89). B. Resulting matrix of transfer entropy. C. Corresponding total TE values (n=83). D. Correlation between ACW-0 and total TE (n=77 that have both rest and task recordings). Correlations were done on median values. Conventions are the same as figure 2

### Total information transfer relates to the steepness of the timescale hierarchy

When we pooled the results from all the different manipulations together, we saw that total information transfer is negatively correlated with the slope of a regression line fitted to the timescale hierarchy: the steeper the hierarchy of INTs, the lower the total TE (figure 7A). This might indicate that in a network with steeper timescale hierarchy and/or higher number of regions following such timescale hierarchy, at every step, more information is downsampled according to the hypothesis we sketched above. Hence, in a network with steeper hierarchy, the relative differences in the information transfer between regions are bigger, resulting in more downsampling. This is exactly what we saw when we correlated slopes of timescale and TE hierarchies in the manipulations of hierarchical synaptic excitation parameter *η* (figure 6D).

**Figure 6.**
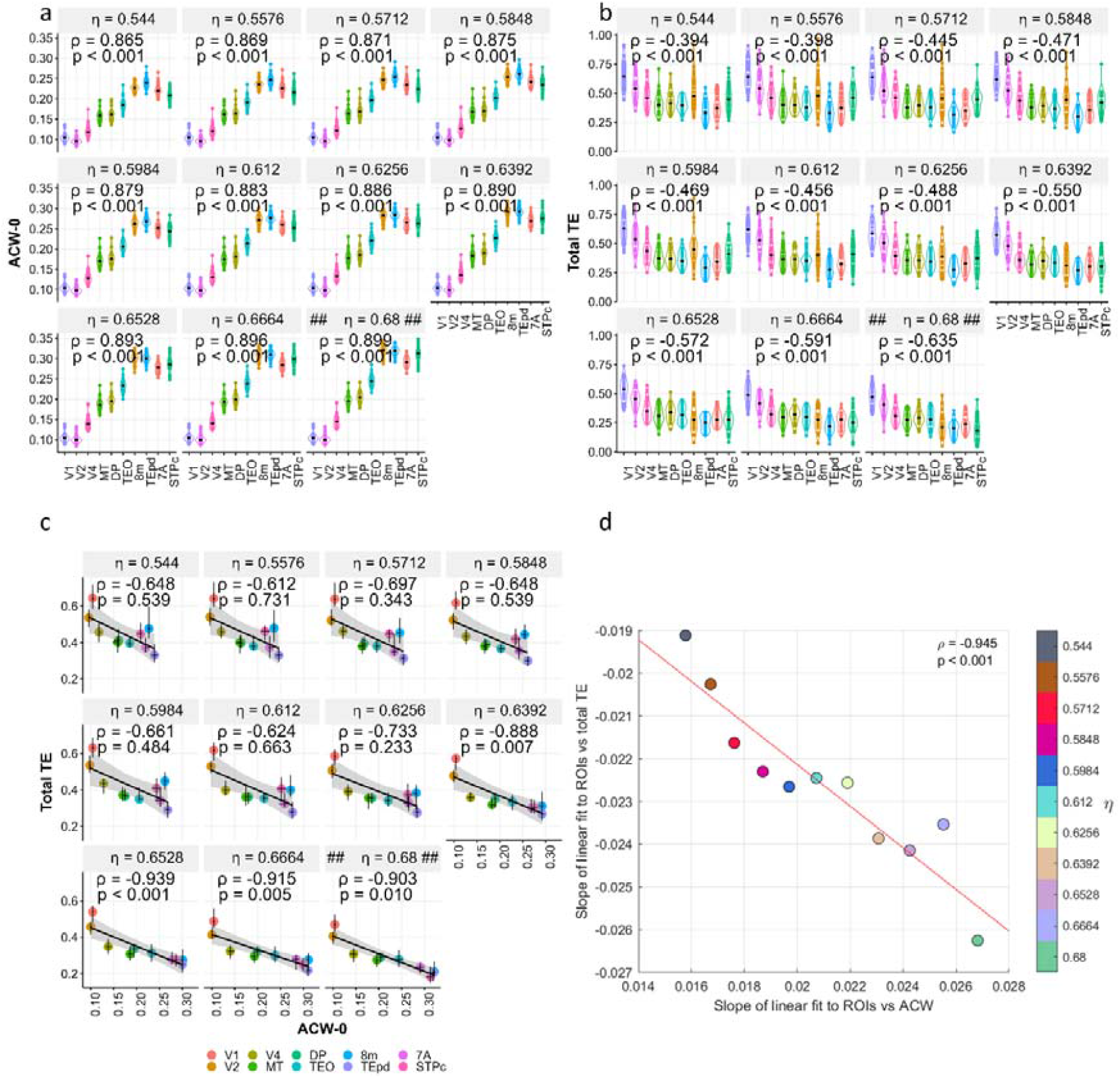
Manipulation of the hierarchy constant. A. ACW-0 values for changing. and p values are results of spearman correlations between ROI orderings and ACW values. B. Total TE (summation of incoming and outgoing) values for changing. Correlations are the same as figure A. C. Correlations between ACW-0 and Total TE for changing. D. The correlation between slope of linear fit to ROIs vs ACWs and same for total TEs. Each dot represents a manipulation in made in the model. In all figures, the panels with ## = 0.68 ## shows the value in figure 2. n = 30 simulations in each a, b and c. The correlations were done on medians of 30 simulations. n = 11 in d.

Interestingly, we saw cluttering of values in the upper and lower extremes of the slopes in figure 7A. To further inspect this, we plotted the histograms of ACW slopes and total TE values. While the topological analysis of the 58-dimensional phase plane of our model is out of the scope of this paper, from a dynamic systems perspective, the bimodal distribution of ACW slopes we saw in figure 7B might indicate a bifurcation^62^: the qualitative change in model dynamics caused by a slight change in one of the parameters. In this context, the few values in the middle might be indicative of a critical regime, right in between hierarchical and non-hierarchical cases. On the other hand, information transfer is distributed more uniformly as we saw in figure 7C. This is in accordance with the decoupling of ACW and TE hierarchies we saw in the manipulation of different model parameters: even in the cases where ACW hierarchy was lost, TE hierarchy usually remained intact but the values increased more or less.

**Figure 7.**
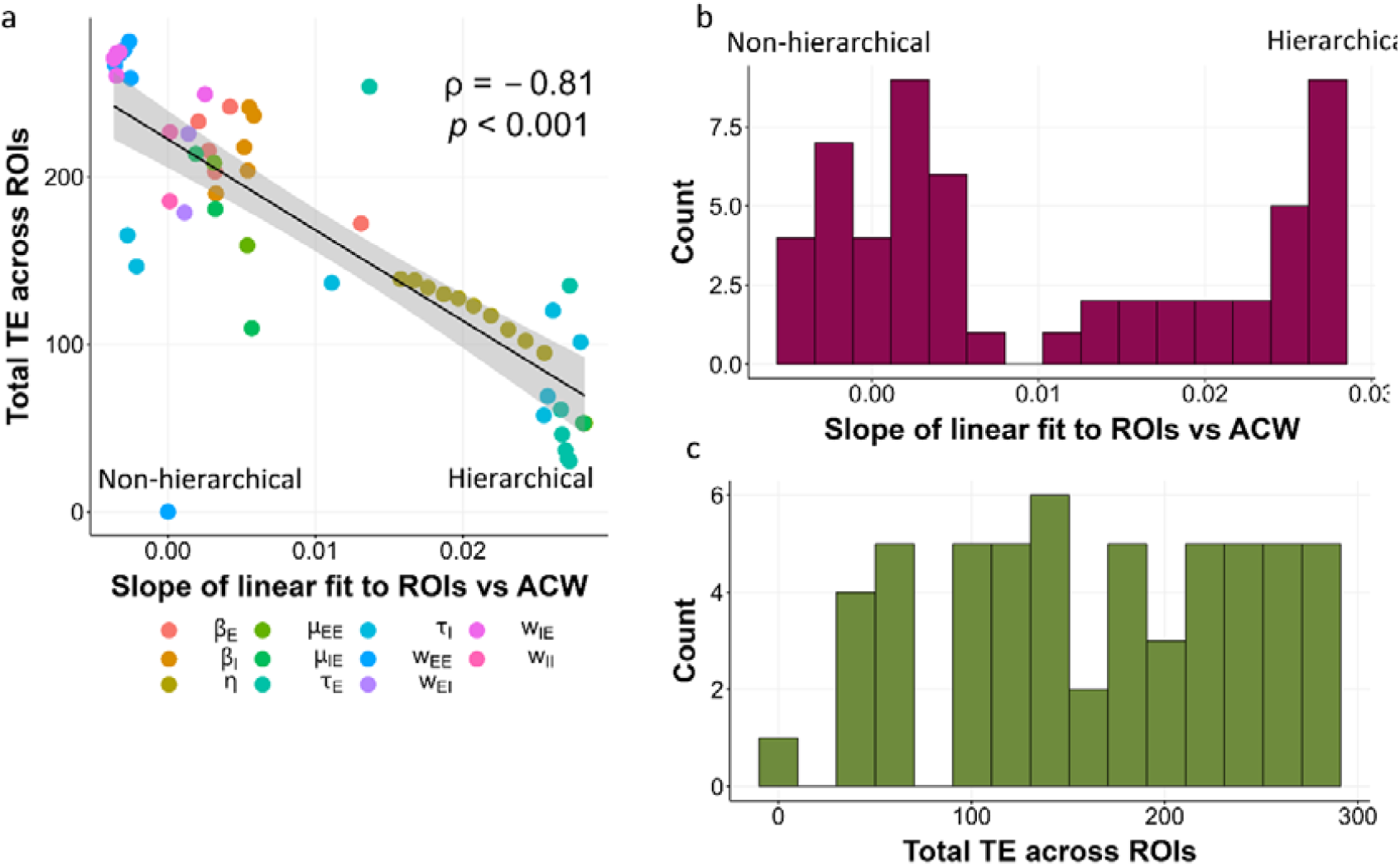
The steepness of timescale hierarchy correlates with the total transfer entropy (TE) across ROIs. A. Spearman correlation between slope of linear fit to ROI orderings and their ACW values and total TE across ROIs in every manipulation we did in the mode (n=56). B. Histogram of slopes showing a bimodal distribution. C. Histogram of total TEs showing a more uniform distribution.

In essence, our results support the hypothesis of INT hierarchy acting as an instrument of gradual downsampling. When we lose the timescale hierarchy in the model, that is, lose the gradual downsampling, this results in an overall increase in total TE in every region, independent of the parameter manipulated. This underscores the key relevance of the timescale hierarchy for information transfer and, as we assume, for the down-sampling of the incoming inputs along the uni-transmodal gradient.

Such down-sampling of information transfer may be altered if not lost in mental disorders. A previous fMRI study by Wengler et al^12^ showed that schizophrenic individuals have lower INT values. We would hypothesize that this would result in more information transfer and thus decreased down-sampling from sensory to association areas which can possibly account for some of the symptoms including certain deficits in sensory processing in schizophrenia.

## LIMITATIONS

While we were able to replicate our modeling findings in the empirical side, we did not do an extensive analysis of task activation within the MEG data since such detailed analyses is beyond the scope of this paper. Instead, we relied on two fMRI studies that utilized the same tasks with the same temporal structure and then applied those regions to our MEG data. We expect the future studies to replicate or disprove our findings on the empirical side not only in MEG and EEG but also in fMRI.

## CONCLUSION

We probed the relationship between the macroscopic uni-transmodal gradient of intrinsic neural timescales (INTs) and information transfer in the brain. Both computational and empirical results show that the hierarchy of INTs with lower INTs at sensory and higher in associative is related to a corresponding hierarchy in information transfer that is higher in sensory and lower in associative regions. Regardless of the parameter that we changed in the model; the resulting loss of the timescale hierarchy always increased the information transfer in every region. Correspondingly, we found that the degree of total information transfer was negatively related to the steepness of timescale hierarchy.

Our findings indicate that one of the functions of the hierarchy of timescales is to modulate information transfer by attenuating it along a uni-transmodal gradient. Such attenuation of the information transfer is well compatible with what is described as ‘down-sampling’ in signal processing literature^40,63^: the brain’s neural activity gradually filters the incoming high-frequency information, along its uni-transmodal timescale hierarchy such that the higher-order regions are predominantly characterized by low-frequency information. This does not only shed a novel light on the role and function of the brain’s timescale hierarchy but also carries major implications for mental disorders like schizophrenia where such attenuation of the inter-regional information transfer might be disrupted.

## Data and Code Availability

The MEG data is available at https://db.humanconnectome.org/. The code to replicate the analyses is at https://github.com/duodenum96/te_acw.

## Statistical Information

We used Wilcoxon tests to compare groups since our data was usually not Gaussian. To estimate error bars, we took 10000 bootstraps on median of the data and estimated 95% confidence intervals. On two-way ANOVAs, we tested the effect of independent variables ROI orederings (1 for V1, 2 for V2 and so on) and the existence or absence of hierarchy (defined as a positive and significant correlation between ROI orderings and ACW-0 values) on the dependent variable total transfer entropy for each ROI. We report the effect of existence of hierarchy in the text. We used the effect size measures difference in location (dil) for Wilcoxon tests and partial *η*^2^ (not to be confused with the parameter *η* in the model) for ANOVA tests.

## Acknowledgements

G.N. is supported by the European Union’s Horizon 2020 Framework Program for Research and Innovation under the Specific Grant Agreement no. 785907 (Human Brain Project SGA2), UMRF, uOBMRI, CIHR, and PSI. He is also grateful to CIHR, NSERC, and SSHRC for supporting the tri-council grant from the Canada-UK Artificial Intelligence (AI) Initiative “The self as agent-environment nexus: crossing disciplinary boundaries to help human selves and anticipate artificial selves” (ES/T01279X/1) (together with Karl J. Friston from the UK).

