## Supplementary Material for "Intrinsic neural timescales attenuate information transfer along the uni-transmodal hierarchy"

**Structural connectivity matrix for the model**

The directed structural connectivity matrix is taken from Markov et al (2014). It is denoted as $FLN$ in the model equations, which corresponds to fraction of labeled neurons. The matrix covers 29 regions in macaque cortex. Inter-areal connection strengths were estimated by counting projecting neurons labeled by a retrograde trace injection and normalizing by dividing this count by the total number of neurons labeled in the injection.

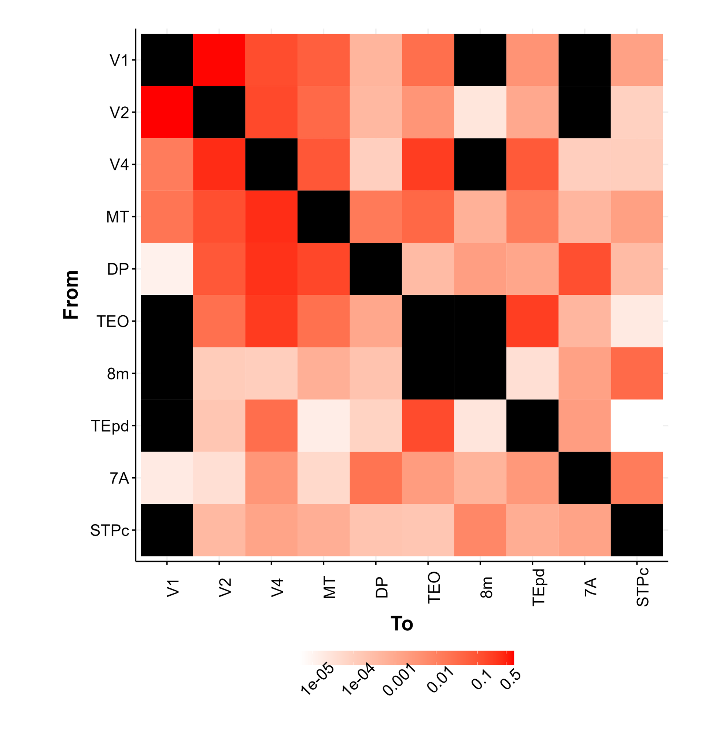

Supplementary Figure 1. Directed connectivity matrix $FLN$.

**Confirmatory analysis for data-driven estimation of Markov lags**

We used a data-driven approach in estimating Markov lags when calculating transfer entropy (see methods). One might argue that this approach biases estimated TE values. To counter this argument, we correlated median of estimated TE values with sum of estimated median of within ($q_{xx}+q_{yy}$) and between ($q_{xy}+q_{yx}$) Markov lags across simulations in the model and subjects in two tasks of the data. None of the correlations were significant. This shows that our TE calculations were not biased by the data-driven estimation method.

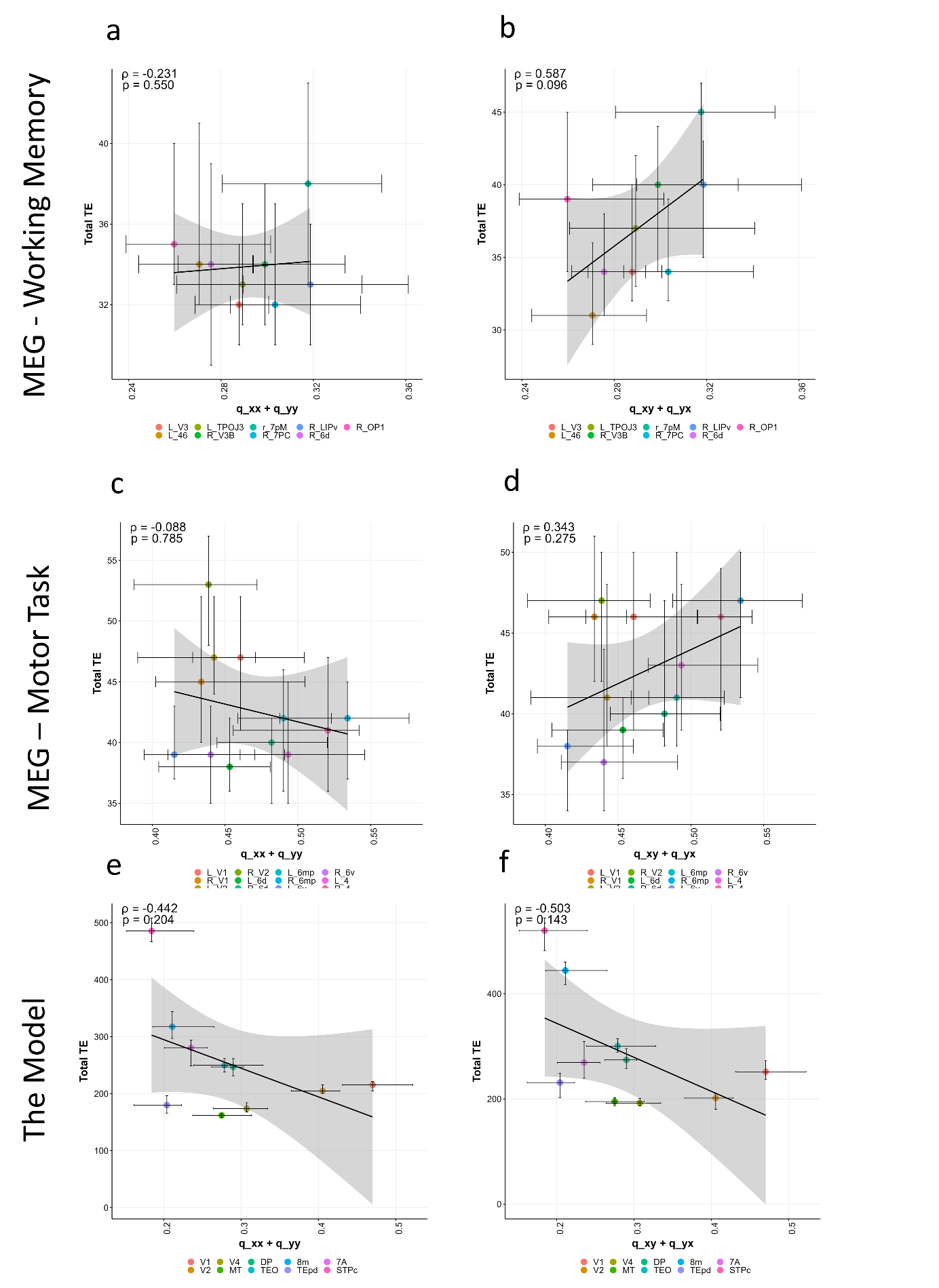

Supplementary Figure 2. Correlation between estimated Markov lags and median TE values. None of the correlations is significant. This shows that estimated TE values are not biased by Markov lag estimation methods.

**Results for the manipulation of other parameters**

Below, we report the results for manipulations of the parameters that we did not report in the main manuscript: $w_{EI}$, $w_{IE}$, $w_{II}$, $\mu_{IE}$, $\beta_{I}$, $\tau_{E}$, $\tau_{I}$. In every figure, panel a shows the violin plots for ACW-0 values per region. Panel b shows the same for total TE (summed incoming and outgoing TE for every region). The statistics of these panels show the correlation between median ACW and TE values and the ordinal variable ROI ordering. In all figures except for $\tau_{E}$, figure c shows the comparison of total TE values in the hierarchical and non-hierarchical network configurations. Hierarchicality was defined as a positive and significant correlation between ROI orderings and ACW-0 values. In every parameter except for $\beta_{I}$, we observed that the total TE is significantly higher in the nonhierarchical case compared to the hierarchical one. The discrepancy in $\beta_{I}$ can be explained by looking at supplementary figure 7 panel A. Our criterion of hierarchy wrongly attributes hierarchy to some networks such as the one in $\beta_{I}$= 0.6318 while it can be observed by looking at the panel A that visually, there is no consistent hierarchy of timescales. For every parameter except $\tau_{E}$, panel D shows the correlation between the median values of ACW-0 and total TE that we saw in panels A and B. This is panel C for $\tau_{E}$ since no network was identified as non-hierarchical in $\tau_{E}$. The TE matrices of all the different model manipulations are given in supplementary figures 10 and 11.

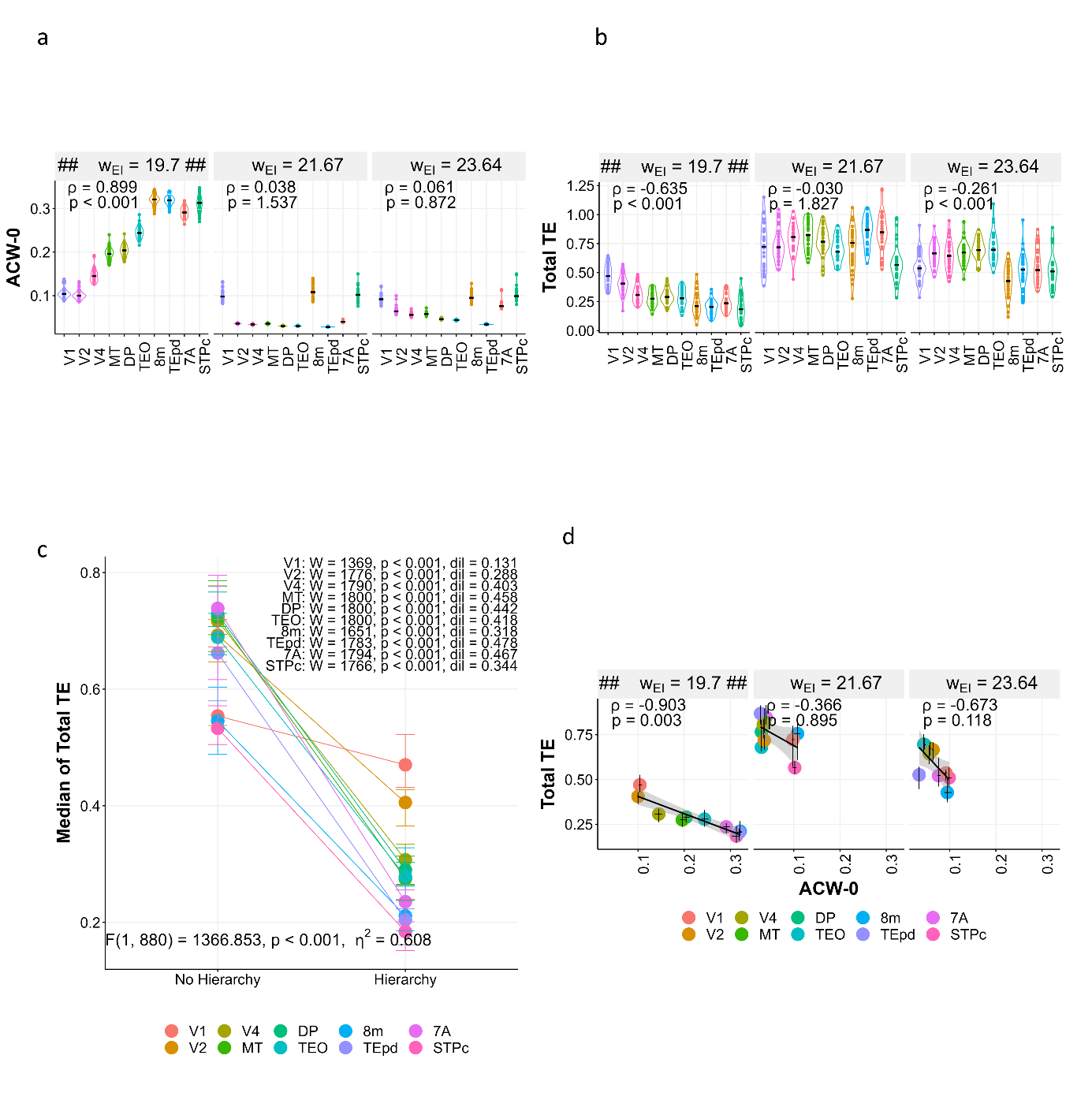

Supplementary Figure 3. Manipulation of local inhibitory-to-excitatory coupling parameter $w_{EI}$. A. ACW-0 values for changing $w_{EI}$. Correlations are the same as figure 2. B. Total TE (summation of incoming and outgoing) values for changing $w_{EI}$. Correlations are the same as figure 2. C. Total TE increases when the hierarchy is lost. We defined the existence of hierarchy as a positive and significant correlation between ROI orderings and ACW-0 values. W: Wilcoxon test statistic. dil: effect size measure difference in location. F value and corresponding statistics p and partial $\eta^{2}$ are results from a two-way ANOVA with variables ROIs and the existence of hierarchy. Values show the results for the dummy variable existence of hierarchy. D. Correlations between ACW-0 and Total TE. In all figures, the panels with ## $w_{EI}$ = 19.7 ## shows the value in figure 2 in the main manuscript.

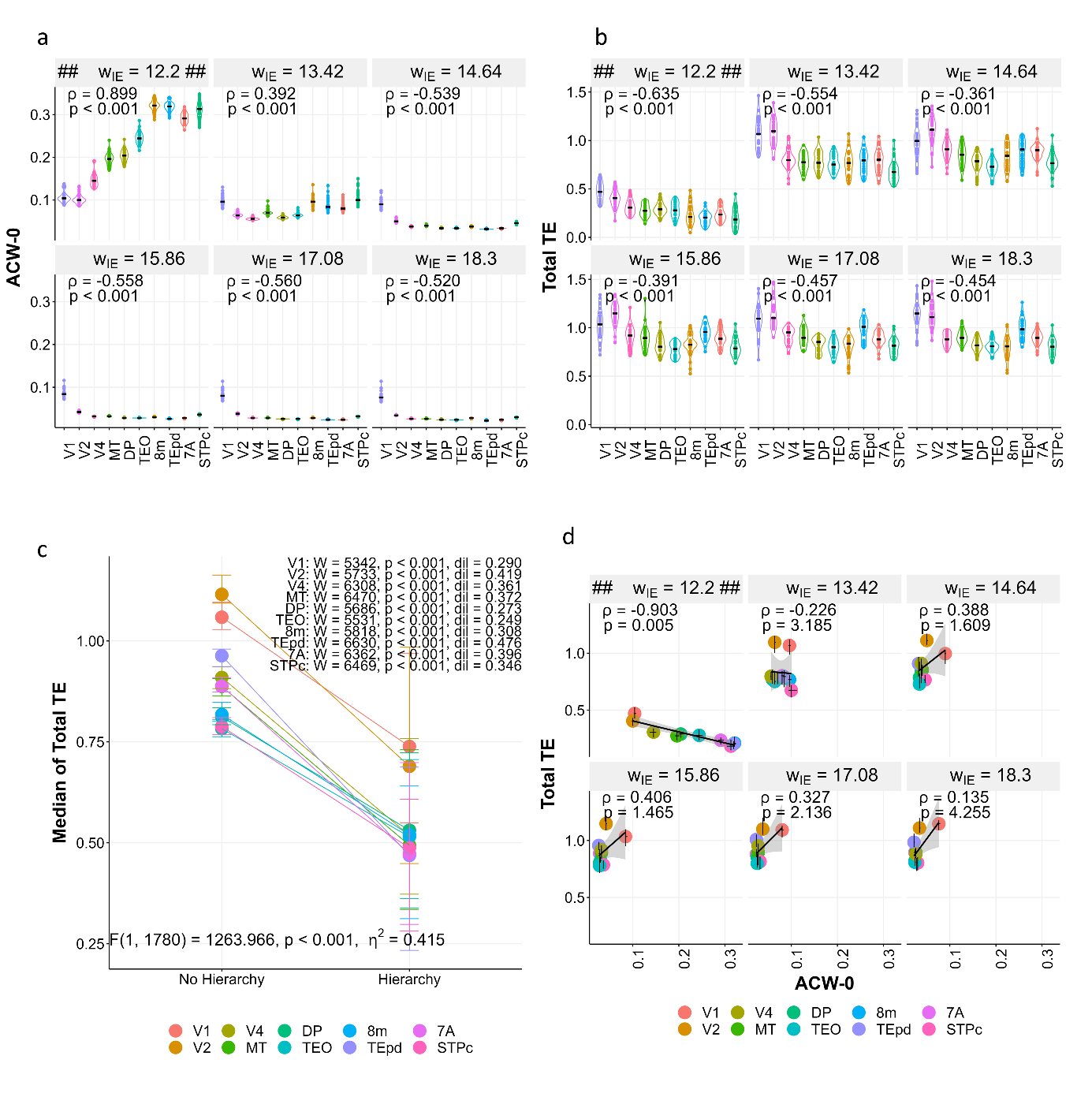

Supplementary Figure 4 Manipulation of local excitatory-to-inhibitory coupling parameter $w_{IE}$. A. ACW-0 values for changing $w_{IE}$. Correlations are the same as figure 2. B. Total TE (summation of incoming and outgoing) values for changing $w_{IE}$. Correlations are the same as figure 2. C. Total TE increases when the hierarchy is lost. We defined the existence of hierarchy as a positive and significant correlation between ROI orderings and ACW-0 values. W: Wilcoxon test statistic. dil: effect size measure difference in location. F value and corresponding statistics p and partial $\eta^{2}$ are results from a two-way ANOVA with variables ROIs and the existence of hierarchy. Values show the results for the dummy variable existence of hierarchy. D. Correlations between ACW-0 and Total TE. In all figures, the panels with ## $w_{IE}$ = 12.2 ## shows the value in figure 2 in the main manuscript.

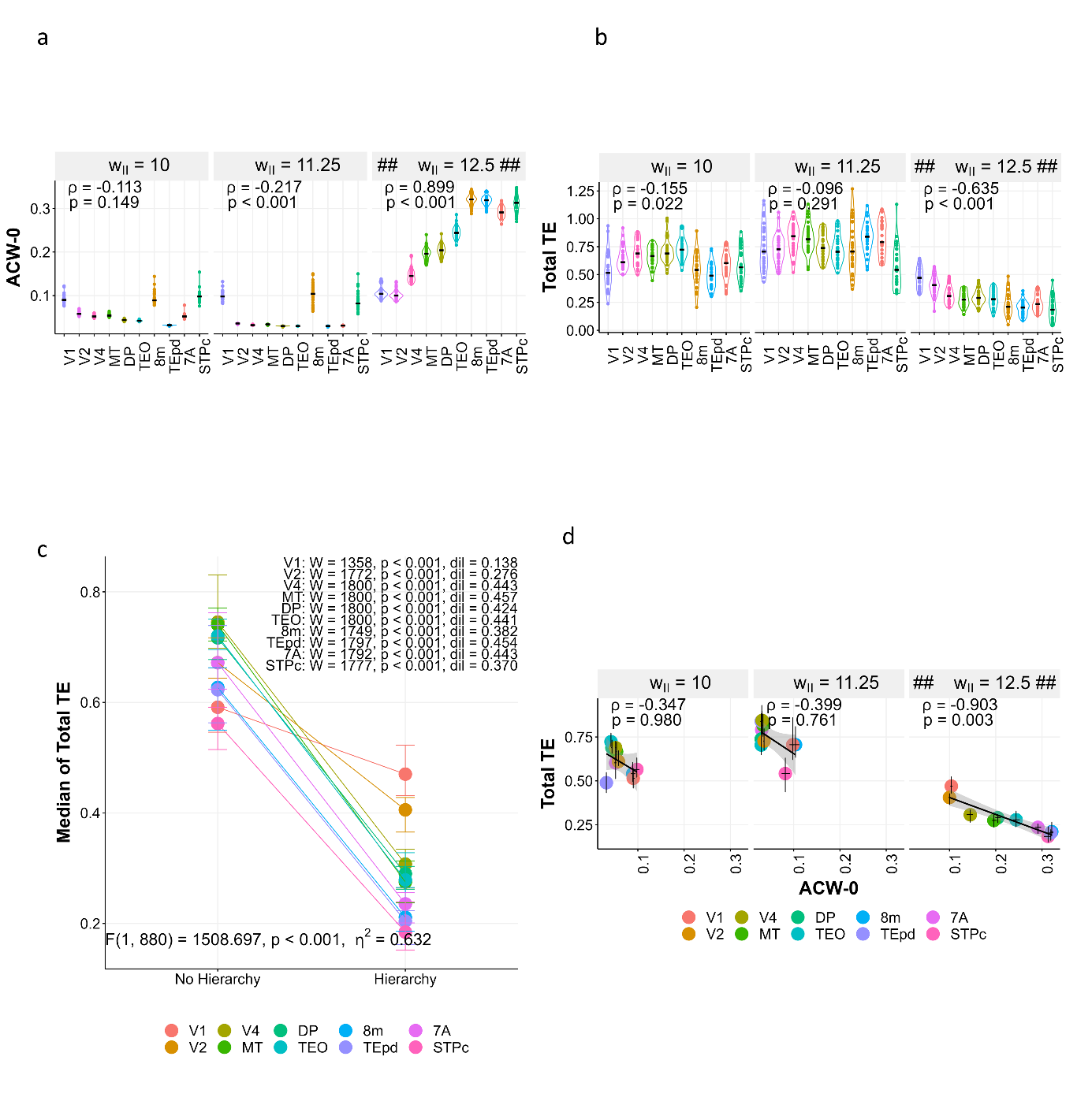

Supplementary Figure 5. Manipulation of local inhibitory-to-inhibitory coupling parameter $w_{II}$. A. ACW-0 values for changing $w_{II}$. Correlations are the same as figure 2. B. Total TE (summation of incoming and outgoing) values for changing $w_{II}$. Correlations are the same as figure 2. C. Total TE increases when the hierarchy is lost. We defined the existence of hierarchy as a positive and significant correlation between ROI orderings and ACW-0 values. W: Wilcoxon test statistic. dil: effect size measure difference in location. F value and corresponding statistics p and partial $\eta^{2}$ are results from a two-way ANOVA with variables ROIs and the existence of hierarchy. Values show the results for the dummy variable existence of hierarchy. D. Correlations between ACW-0 and Total TE. In all figures, the panels with ## $w_{II}$ = 12.5 ## shows the value in figure 2 in the main manuscript.

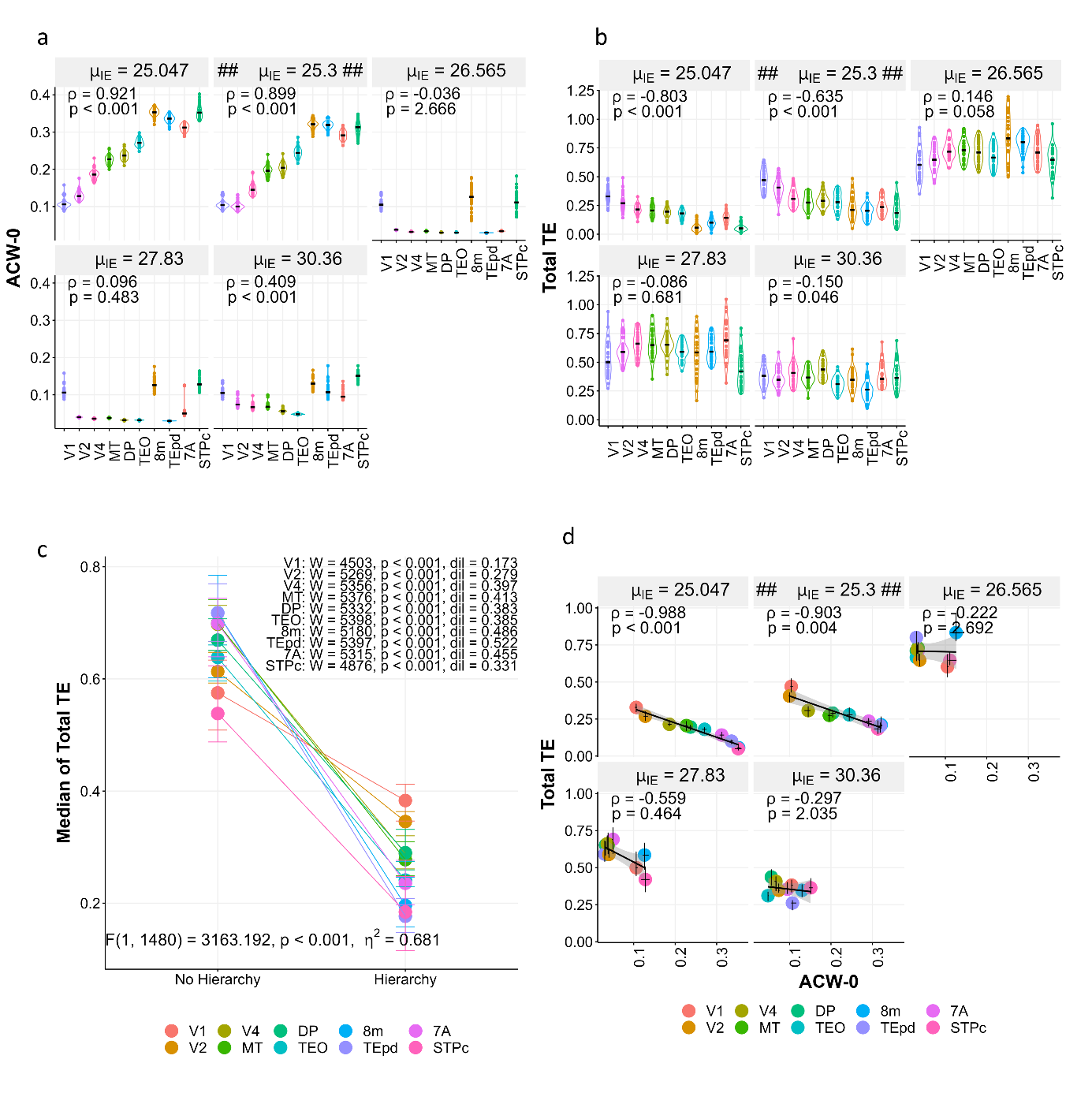

Supplementary Figure 6. Manipulation of global excitatory-to-inhibitory coupling parameter $\mu_{IE}$. A. ACW-0 values for changing $\mu_{IE}$. Correlations are the same as figure 2. B. Total TE (summation of incoming and outgoing) values for changing $\mu_{IE}$. Correlations are the same as figure 2. C. Total TE increases when the hierarchy is lost. We defined the existence of hierarchy as a positive and significant correlation between ROI orderings and ACW-0 values. W: Wilcoxon test statistic. dil: effect size measure difference in location. F value and corresponding statistics p and partial $\eta^{2}$ are results from a two-way ANOVA with variables ROIs and the existence of hierarchy. Values show the results for the dummy variable existence of hierarchy. D. Correlations between ACW-0 and Total TE. In all figures, the panels with ## $\mu_{IE}$ = 25.3 ## shows the value in figure 2 in the main manuscript in the main manuscript.

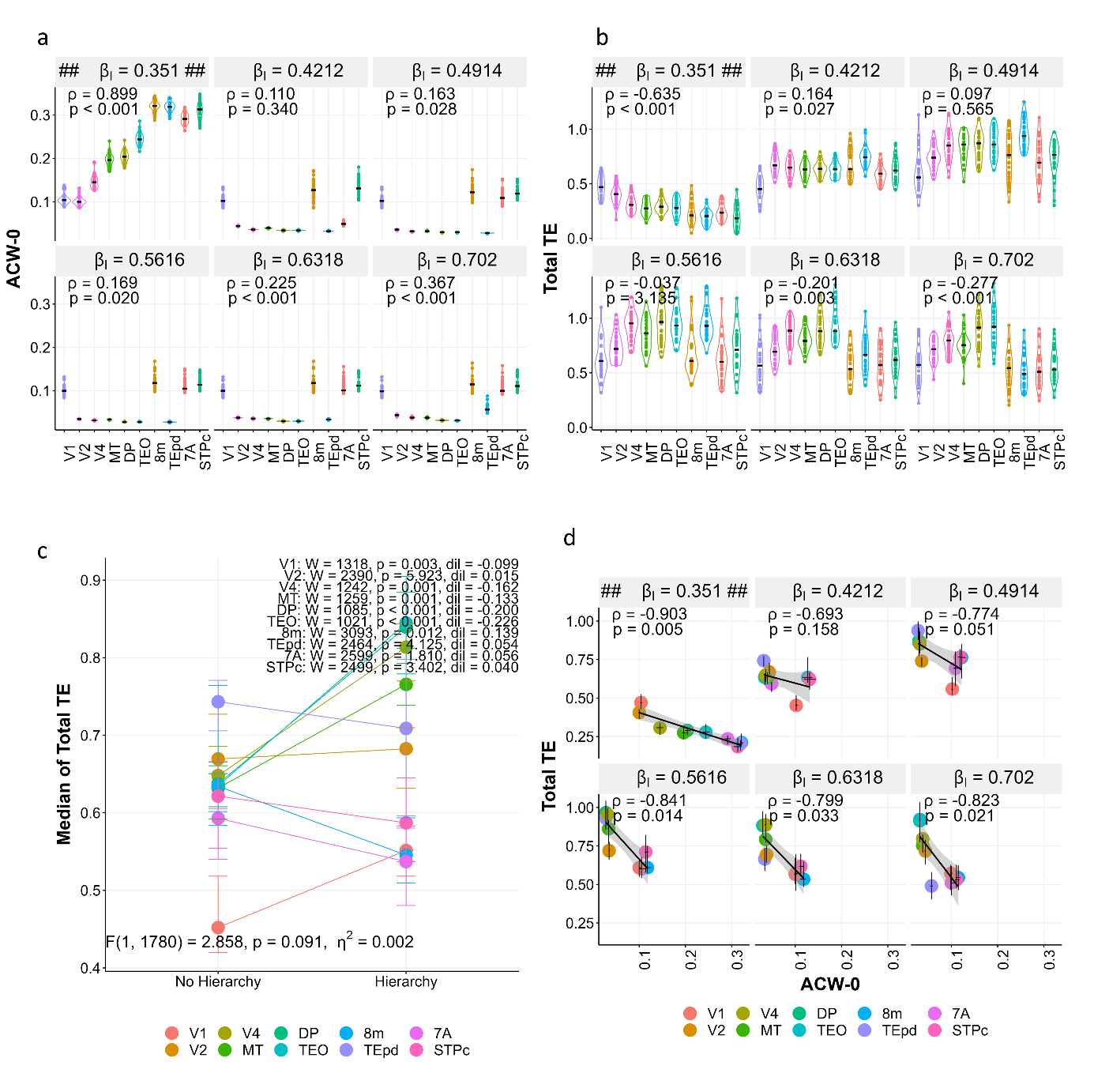

Supplementary Figure 7. Manipulation of slope of the rectifier curve $\beta_{I}$. A. ACW-0 values for changing $\beta_{I}$. Correlations are the same as figure 2. B. Total TE (summation of incoming and outgoing) values for changing $\beta_{I}$. Correlations are the same as figure 2. C. Total TE increases when the hierarchy is lost. We defined the existence of hierarchy as a positive and significant correlation between ROI orderings and ACW-0 values. W: Wilcoxon test statistic. dil: effect size measure difference in location. F value and corresponding statistics p and partial $\eta^{2}$ are results from a two-way ANOVA with variables ROIs and the existence of hierarchy. Values show the results for the dummy variable existence of hierarchy. D. Correlations between ACW-0 and Total TE. In all figures, the panels with ## $\beta_{I}$ = 0.351 ## shows the value in figure 2 in the main manuscript.

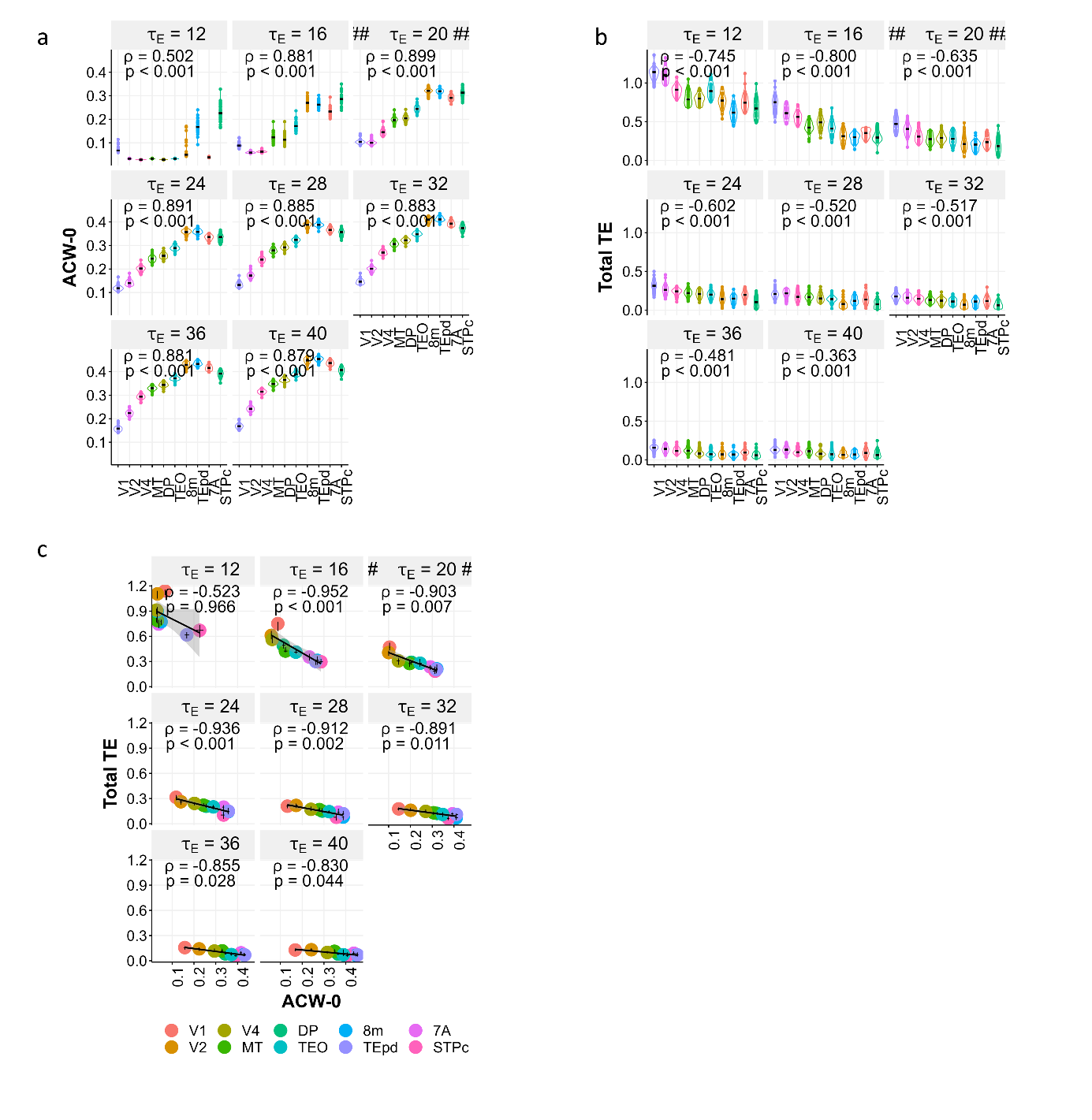

Supplementary Figure 8. Manipulation of time constant for excitatory populations $\tau_{E}$. A. ACW-0 values for changing $\tau_{E}$. Correlations are the same as figure 2. B. Total TE (summation of incoming and outgoing) values for changing $\tau_{E}$. Correlations are the same as figure 2. C. Correlations between ACW-0 and Total TE. In all figures, the panels with ## $\tau_{E}$ = 20 ## shows the value in figure 2 in the main manuscript.

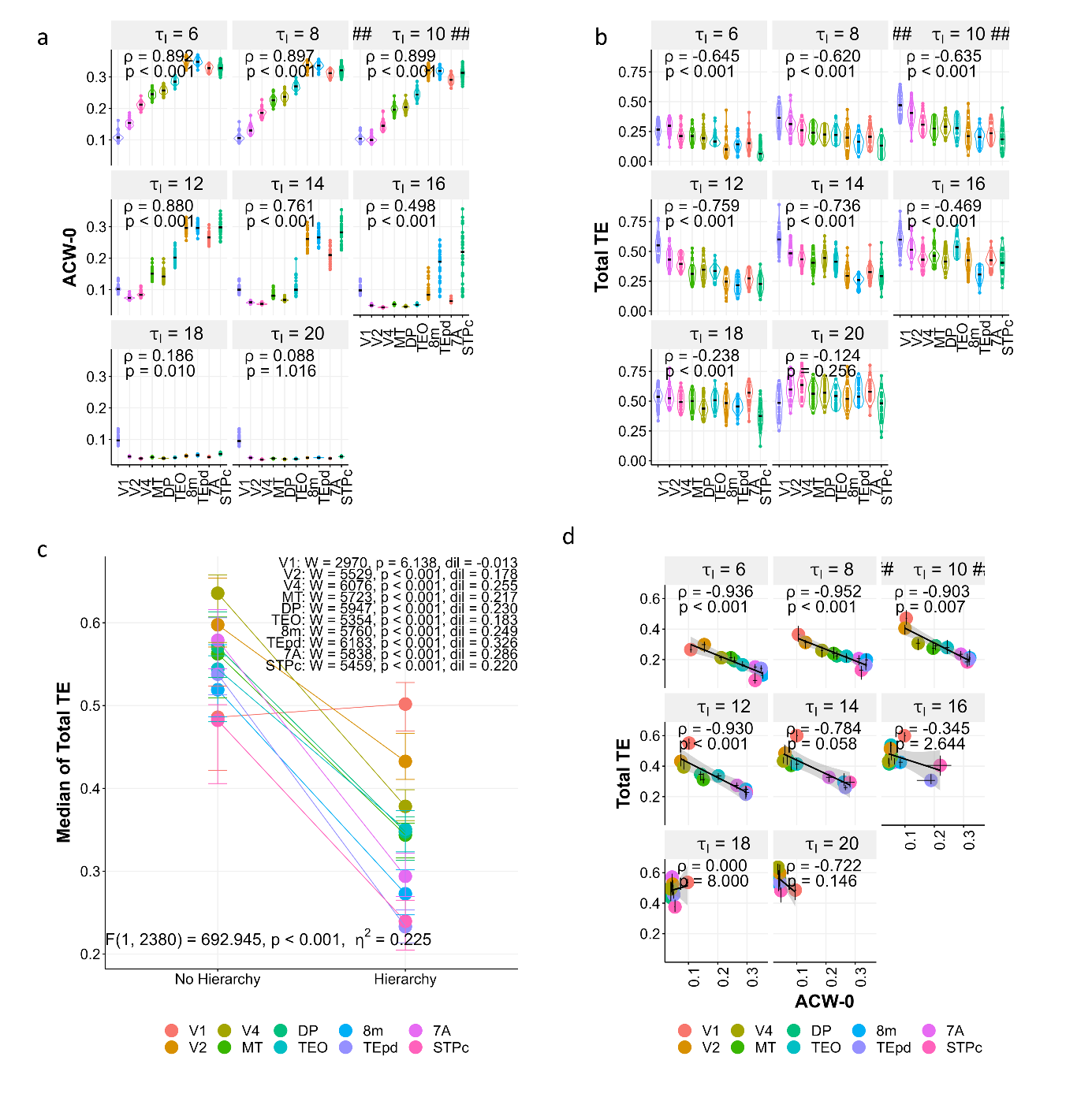

Supplementary Figure 9. Manipulation of the time constant for inhibitory population $\tau_{I}$. A. ACW-0 values for changing $w_{EE}$. Correlations are the same as figure 2. B. Total TE (summation of incoming and outgoing) values for changing $\tau_{I}$. Correlations are the same as figure 2. C. Total TE increases when the hierarchy is lost. We defined the existence of hierarchy as a positive and significant correlation between ROI orderings and ACW-0 values. W: Wilcoxon test statistic. dil: effect size measure difference in location. F value and corresponding statistics p and partial $\eta^{2}$ are results from a two-way ANOVA with variables ROIs and the existence of hierarchy. Values show the results for the dummy variable existence of hierarchy. D. Correlations between ACW-0 and Total TE. In all figures, the panels with ## $\tau_{I}$ = 10 ## shows the value in figure 2 in the main manuscript.

**Transfer Entropy Matrices in the Model Manipulations**

Below, the transfer entropy matrices for all different model manipulations are given (averaged across simulations).

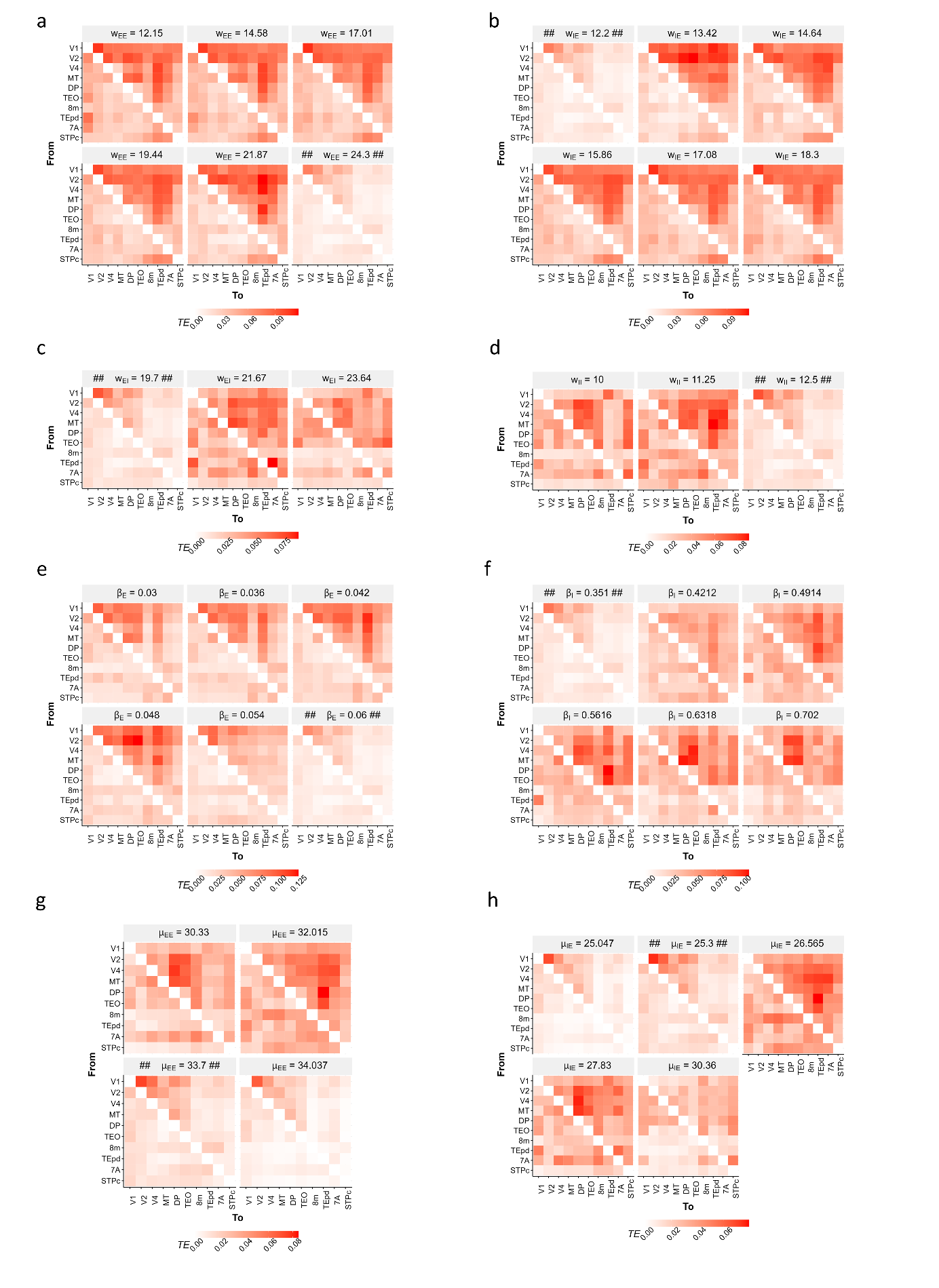

Supplementary Figure 10. Transfer entropy matrices for model manipulations (continued in supplementary figure 11).

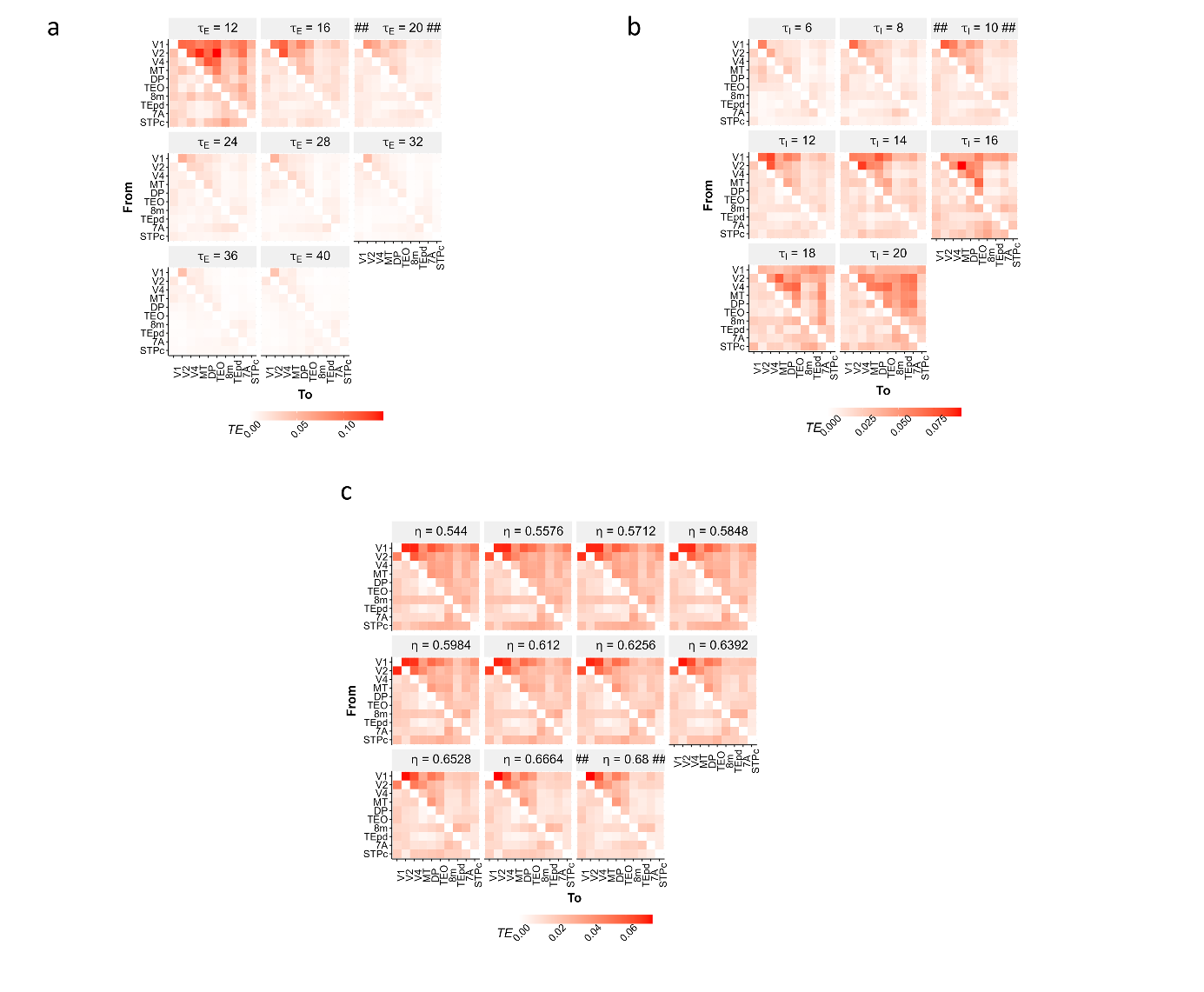

Supplementary Figure 11. Transfer entropy matrices for model manipulations.

**Scatterplots of** $\boldsymbol{\eta}$**s with Spearman** $\boldsymbol{\rho}$ **values of correlation between ROI orderings and ACW / TE values**

Below, we show the scatterplots between spearman correlation $\rho$ values between ROI orderings and ACW / TE values with increasing $\eta$.

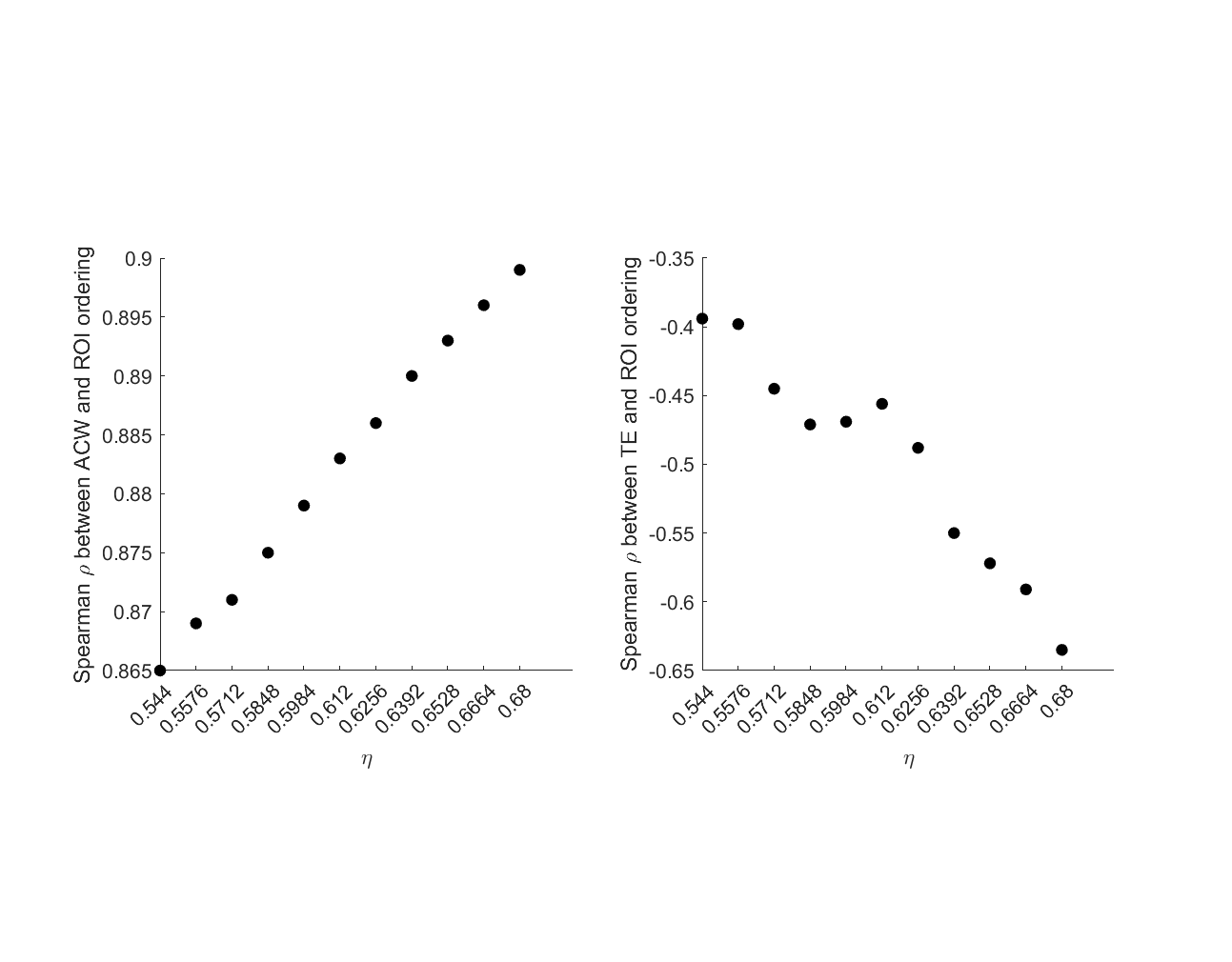

Supplementary Figure 12. Scatterplots of ηs with Spearman ρ values of correlation between ROI orderings and ACW / TE values

**The effect of shuffling of ROI orders in empirical data**

To show that our way of estimating ROI orderings is does not give random results, we randomized the order of ROIs 10000 times and fit linear regression curves between ROI orderings and median total TE values. This created null distributions for us. Our ACW-0 based linear regression slope had a 2-sided p-value (calculated as the probability of getting as extreme or more extreme slope in the null distribution of randomized ROI orderings) was 0.0074 (supplementary figure 12) for working memory and 0.0764 for the motor task.

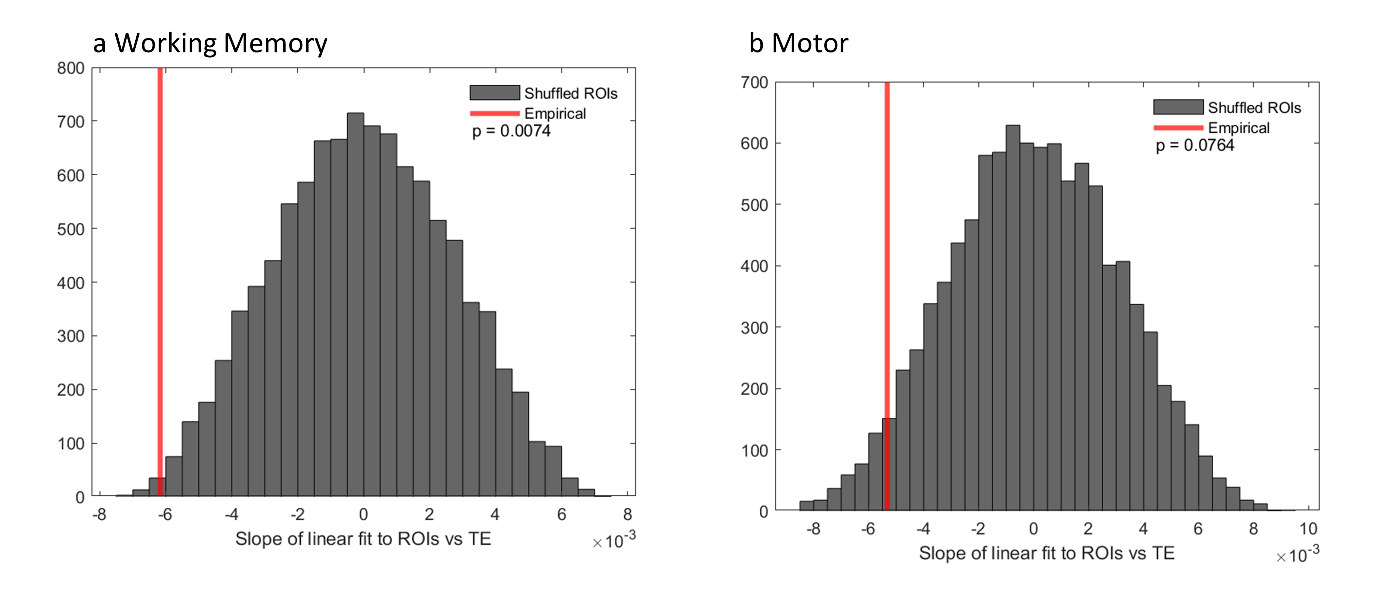

Supplementary Figure 13. To counter the argument that ROI orderings in empirical data can be achieved randomly, we randomized the ROI orderings 10000 times and estimated the slope of TE hierarchy in each ROI order. The histograms show the counts of the slopes. Two-sided p values shown are calculated as seeing a slope as extreme or more extreme as our empirical ROI ordering.

**Results for the motor task**

For the motor task, we ordered the regions according to their ACW-0 values at rest (supplementary figure 13A). As expected, this showed a significant and positive Spearman correlation ($\rho$ = 0.431, p < 0.001) between the ROI orderings and their ACW values. The resulting TE matrix can be seen in supplementary figure 13B. The total transfer entropy values for the regions followed the ACW hierarchy and were consistent with the simulation results. While it was not as marked as in the simulations, we saw a significant negative correlation between the ACW-based ROI orderings and median total TE values (rho = -0.129, p < 0.001) (figure 8C). To show that our way of estimating ROI orderings is does not give random results, we randomized the order of ROIs 10000 times and fit linear regression curves between ROI orderings and median total TE values. This created null distributions for us. Our ACW-0 based linear regression slope had a 2-sided p-value (calculated as the probability of getting as extreme or more extreme slope in the null distribution of randomized ROI orderings) was 0.0764 (the histogram can be seen in supplementary figure 12). Similar to our models, median of total TE values per ROI correlated negatively with their median ACWs: the longer the ACW of a region, the lower its total TE, e.g., its information flow (rho = -0.66, p = 0.024, figure 8D).
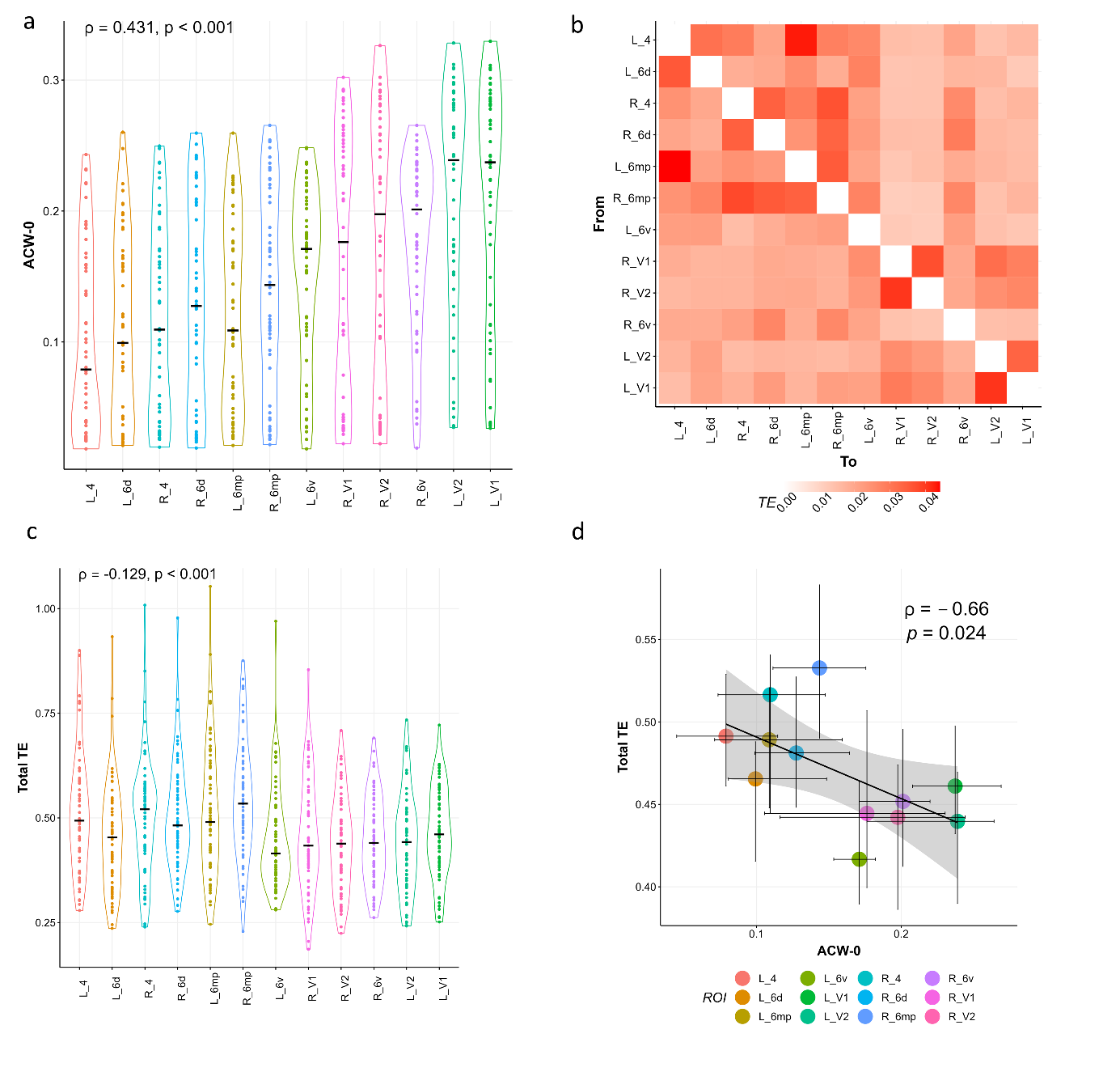

Supplementary Figure 14. Replication of results in the HCP-MEG dataset motor task. A. Resting state ACW-0 values, ordered from lowest to highest (n = 89). B. Resulting matrix of transfer entropy. C. Corresponding total TE values (n = 61). D. Correlation between ACW-0 and total TE (n = 56 that have both rest and task scans). Conventions are the same as figure 2 in the main manuscript. Correlations were done on median values).
